# GLM-Prior: a genomic language model for transferable sequence-derived priors in gene regulatory network inference

**DOI:** 10.1101/2025.06.29.662198

**Authors:** Claudia Skok Gibbs, Angelica Chen, Richard Bonneau, Kyunghyun Cho

## Abstract

Gene regulatory network (GRN) inference depends on high-quality prior knowledge, yet curated priors are often incomplete or unavailable across species and cell types. We present GLM-Prior, a genomic language model fine-tuned to predict transcription factor (TF)-target gene interactions from nucleotide sequence, and benchmark it as a sequence-derived strategy for constructing TF-gene prior matrices. We integrate GLM-Prior with PMF-GRN, a probabilistic matrix factorization model, to create a dual-stage pipeline that combines sequence-derived priors with single-cell gene expression data for prior-conditioned GRN inference. Across six human, mouse, and yeast cell-line contexts, GLM-Prior performance scales with positive label abundance and TF coverage in training data, and shows above-chance agreement with independent reference networks in well-annotated mammalian contexts. We evaluate single-species, species-transfer, and multi-species training paradigms, finding that GLM-Prior can construct informative priors across related mammalian species, while transfer to yeast remains near chance. Comparisons with accessibility-based priors across multiple GRN inference methods show that GLM-Prior achieves the highest prior performance in four of five mammalian cell lines. Under these benchmarks, prior quality largely constrains achievable GRN inference performance, with expression-based inference providing prior-dependent refinement. Together, these results position GLM-Prior as a benchmarked workflow for transferable, sequence-derived prior construction in systems where matched experimental assays are unavailable.

## 1 Introduction

Gene regulatory networks (GRNs) map the transcriptional relationships between transcription factors (TFs) and their target genes, providing a framework for understanding cellular function and gene expression control in cells (1; 2). Constructing useful GRN models requires integrating large-scale sequencing datasets, such as single-cell RNA-seq and ATAC-seq, to infer regulatory interactions that cannot be measured experimentally (3). However, the accuracy of GRN inference depends heavily on the quality of the prior knowledge used to guide the inference algorithm (4; 5). In the context of GRNs, prior knowledge refers to an initial matrix of putative TF-gene regulatory interactions, which provides structural guidance to the inference algorithm by defining which TF-gene pairs are more likely to represent regulatory edges. Recent methods have demonstrated that better prior knowledge, which captures biologically meaningful regulatory signals, improves agreement with reference networks and supports more informative GRN inference, highlighting the critical role of prior knowledge in improving GRN inference quality (6; 7; 8; 9).

Prior knowledge for well-studied species is typically constructed using databases of experimentally validated TF-gene interactions (10; 11). For less-characterized organisms, prior knowledge is inferred by combining structural genomic data with sequencing information (12). The Inferelator 3.0’s Inferelator-Prior performs TF motif enrichment within open chromatin regions to define edges between proximally bound TFs to target genes (6). CellOracle’s base GRN draws edges between TFs bound within promoter regions of accessible target genes (8). SCENIC’s CisTarget constructs prior knowledge by ranking enriched motifs in accessible regions using genome-wide motif scores and a hidden Markov model to predict TF-gene interactions (9). While these methods demonstrate the importance of prior knowledge in GRN inference, they also expose key limitations. Motif annotation remains incomplete (13), chromatin accessibility data is noisy and cell-type specific (14), and existing methods often fail to capture long-range regulatory interactions (15). Further, these existing approaches cannot create prior knowledge matrices which generalize to other species. These limitations underscore the need for benchmarked approaches to constructing transferable prior knowledge matrices, particularly for poorly characterized organisms (4; 16).

A promising direction is to incorporate transformer-based genomic foundation models as components of prior construction workflows. Architectures such as the Nucleotide Transformer (17) have demonstrated strong performance across genomic prediction tasks, including chromatin accessibility, gene expression, and transcription factor binding (18; 19; 20; 21). These models can learn sequence-associated features across large genomic corpora, making them attractive candidates for constructing TF-gene priors when matched regulatory assays are unavailable. When trained or fine-tuned across multiple species, genomic language models may preserve predictive signal across related organisms, supporting a transfer-learning paradigm in which models trained in well-annotated species can be evaluated in less-characterized systems. In this way, genomic language models offer a potential route to expand prior knowledge construction beyond organisms and cell types with existing curated interaction data, while requiring careful benchmarking against available reference networks.

In this work, we present **GLM-Prior**, a sequence-based workflow for constructing TF-gene prior knowledge matrices to support downstream GRN inference. GLM-Prior is built by fine-tuning the 250 million parameter Nucleotide Transformer (17), pretrained on the genomes of 850 species, into a sequence classification model for predicting TF-gene regulatory labels from available interaction databases. Unlike motif-based approaches that rely on the proximity of TF motifs to accessible promoter regions, GLM-Prior uses paired nucleotide sequences consisting of TF binding motifs and gene bodies. These sequences are jointly encoded by the transformer and passed through a classification head, which outputs a score for each TF-gene pair. To construct a binary prior knowledge matrix suitable for downstream GRN inference, we apply an optimized classification threshold that converts these scores into putative positive and negative prior edges. Since most TF-gene pairs are unlabeled and the set of true negatives is incomplete, the model is trained using class-weighted loss and balanced sampling to mitigate class imbalance during training. We evaluate whether this sequence-derived prior construction strategy improves agreement with held-out reference networks across species and cell-type contexts.

We investigate the generalizability of GLM-Prior by evaluating three training paradigms: (1) single-species models trained and evaluated on the same organism; (2) transfer learning, in which models trained in one species are used to predict regulatory interactions in an unseen species; and (3) multi-species training, where a single model is fine-tuned on data from multiple organisms. Across these settings, performance scales with abundance and diversity of positive class labels and TF coverage in the training data. Human and mouse cell lines with rich label sets yield higher AUPRCs than yeast, where extreme class imbalance and sparse labels limit generalization. Transfer learning between evolutionarily related species (human and mouse) maintains or modestly improves performance relative to single-species models, suggesting that GLM-Prior preserves sequence-associated predictive signal across closely related mammalian species. In contrast, transfer learning between mammals and yeast does not exceed near-chance performance, indicating that this transfer signal does not extend uniformly to more evolutionarily distant organisms under the current benchmark. Multi-species training on human, mouse, and yeast preserves accuracy comparable to the best single-species or transfer models in most cell lines, with only mild trade-offs in some human contexts, and produces a sequence-derived prior with comparable benchmark performance across evaluated contexts, without requiring a separately trained model for each target species.

To evaluate the utility of GLM-Prior, we incorporate it into a dual-stage GRN inference pipeline. In the first stage, GLM-Prior generates a prior knowledge matrix of putative TF-gene interactions from nucleotide sequence. In the second stage, this matrix is used to constrain PMF-GRN (7), a probabilistic matrix factorization model that infers GRNs from single-cell gene expression data. This framework allows us to isolate and compare the contributions of prior construction and expression-based inference to benchmarked GRN performance against available reference networks.

Building on this, we benchmark GLM-Prior against accessibility-based priors (Inferelator-Prior (6), and CellOracle’s baseGRN (8)), and integrate each prior with it’s downstream GRN inference algorithm (PMF-GRN (7), the Inferelator 3.0 (6), and CellOracle (8)) across yeast, mouse, and human cell lines. GLM-Prior performs at or above chance across the six evaluated contexts, with the clearest gains in mammalian settings with stronger label coverage. In contrast, accessibility-based priors remain competitive in settings dominated by proximal regulatory logic, such as yeast, but show more variable agreement with the corresponding mammalian reference networks. When these priors are passed into downstream GRN inference, GRN inference typically yields modest gains when the prior is already strong, and larger improvements are observed only when the prior is weak or inaccurate. Critically, the identity of the best-performing GRN in each cell line almost always matches the identity of the best prior, and no GRN inference method is uniformly superior across priors or species. Together, these results indicate that prior construction, rather than the choice of GRN inference algorithm, is the primary driver of benchmarked GRN performance.

Together, our findings position GLM-Prior as a benchmarked workflow for transferable, sequence-derived prior construction in GRN inference. Across the evaluated contexts, we assess how well sequence-derived priors align with available reference networks and how those priors shape downstream expression-based inference. Prior quality strongly influences benchmarked GRN performance, while downstream GRN methods primarily refine or contextualize the structure supplied by the prior. In this work, we characterize GLM-Prior’s generalization across species and cell-type settings, compare it with accessibility-based prior construction strategies, and dissect how different GRN inference algorithms interact with these priors.

## Results

### Dual-stage training pipeline overview

We develop a dual-stage training pipeline to evaluate sequence-derived prior construction and prior-conditioned GRN inference by integrating GLM-Prior with probabilistic modeling of single-cell expression data (Figure 1). The pipeline is designed to decouple the construction of prior knowledge from the expression-driven GRN inference step, allowing each component to be independently optimized and evaluated. In this framework, the first stage generates a sequence-informed prior knowledge matrix using a fine-tuned genomic language model (GLM-Prior), while the second stage uses this matrix as structural input to the probabilistic matrix factorization model (PMF-GRN (7)) to infer a GRN.

**Figure 1:**
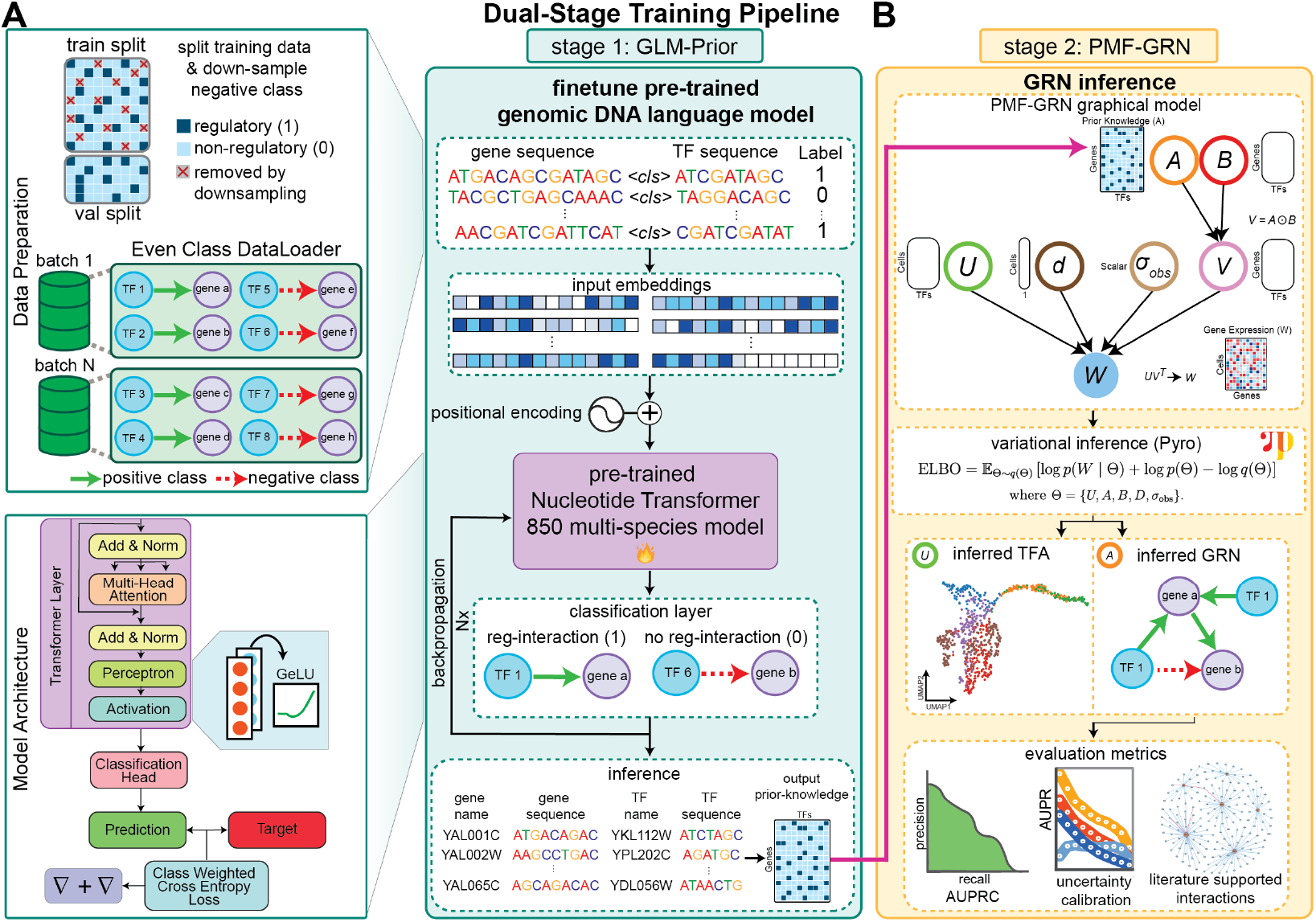
Schematic of the dual-stage training pipeline. (**A**) GLM-Prior is trained on concatenated TF binding motifs and gene body sequences labeled as annotated positive interactions or unlabeled negative examples. Input data is split into training and validation sets, where the training set is downsampled and placed into even class batches using a custom DataLoader. These input sequences are embedded and encoded by the transformer architecture, and passed through a classification head to predict the probability of a positive regulatory label. After training, inference is performed on all TF-gene pairs to generate a prior knowledge matrix. (**B**) Inferred prior knowledge is passed to PMF-GRN to provide a structural constraint to the probabilistic graphical model during inference. PMF-GRN infers transcription factor activity and a GRN for the input cell-line dataset. The resulting GRN is then evaluated using AUPRC and uncertainty calibration with an independent reference network.

### Stage 1: Constructing sequence-informed prior knowledge with GLM-Prior

In the first stage of the pipeline, we construct a prior knowledge edge matrix representing putative positive regulatory (1) and unlabeled or negative (0) entries between TFs and target genes (Figure 1A). To generate these predictions, we develop GLM-Prior, a transformer-based sequence classification model built by fine-tuning the 250 million parameter Nucleotide Transformer (17) model, which was pretrained on the genomes of 850 different species. The primary goal of GLM-Prior is to use sequence information to score TF-gene pairs and construct an informative prior knowledge matrix for downstream GRN inference.

GLM-Prior takes as input paired nucleotide sequences consisting of TF binding motifs and gene body regions, along with labels representing annotated positive interactions and unlabeled negative examples. These sequences are concatenated and encoded into a shared representation using the transformer’s embedding and encoding layers. This representation is passed through a classification layer that outputs the probability of a positive regulatory label. During training, the model evaluates a range of classification thresholds, selecting the threshold that maximizes the F1 score on the validation set. This optimal threshold is used to binarize the predicted probabilities into positive and negative prior entries for the final prior knowledge matrix.

Since GRN datasets are often highly imbalanced, with far fewer positive interactions compared to negative interactions, we apply several strategies to mitigate this imbalance during training. Specifically, we apply a downsampling rate to the negative class and use a class-weighted cross-entropy loss, tuning the negative class weight through hyperparameter search. Additionally, to further ensure stable training, we implement a custom DataLoader to create training batches with equal numbers of positive and negative examples, ensuring balanced gradient updates.

After training, GLM-Prior performs inference across all provided TF-gene pairs, generating a dense probability prior knowledge matrix. The predicted probabilities are then binarized using the optimal F1 score-derived classification threshold, generating a binary prior knowledge matrix of putative regulatory edges for a given organism. Although we can use the probabilities directly, we binarize the prior knowledge matrix to align with existing methods and enable more direct comparison. We then either use this sequence-based prior against an indendent reference network or pass it into the second stage of the pipeline, enabling modular information transfer from sequence-derived prior construction to expression-based GRN inference.

### Stage 2: Refining GRN inference from single-cell data with PMF-GRN

In the second stage of the pipeline, the prior knowledge matrix predicted by GLM-Prior serves as a structured input to our previously developed model, PMF-GRN (7) (Figure 1B). PMF-GRN leverages this sequence-informed prior knowledge to enhance the inference of GRNs from single-cell gene expression data. It does so by using probabilistic matrix factorization, which decomposes the observed gene expression into latent factors representing transcription factor activity (TFA) and TF-target gene regulatory relationships.

Concretely, PMF-GRN models the expression matrix *W* using the likelihood function *p*(*W*|*U, V* = *A* ⊙ *B, σ*_*obs*_, *d*), where 𝔼 [*W*] ≈ *d* ⊙ *UV* ^⊤^, with per-cell sequencing depth *d* and observation noise *σ*_*obs*_. Here, *U* is a nonnegative cells by TFs matrix of TF activities, and *V* is a genes by TFs matrix, which is further factorized as *V* = *A* ⊙ *B. A* ∈ (0, 1) is defined as the degree of existence of a TF-target gene interaction and *B* ∈ ℝ is its signed effect. Each latent variable, *U*, *A, B, d*, and, *σ*_*obs*_ is assumed independent *a priori*. In this formulation, the GLM-Prior predicted prior knowledge matrix provides the hyperparameters for the logistic-normal prior on *A*, anchoring factors to specific TFs and mitigating column-label identifiability inherent to matrix factorization.

Using variational inference, PMF-GRN estimates the posterior distribution of all latent variables. The posterior mean of *A* defines the inferred GRN, while the posterior variance provides an edge-level uncertainty score. These uncertainty estimates enable confidence-weighted interpretation of TF-gene edges and are especially valuable when reference regulatory annotations are incomplete or unavailable.

The combination of GLM-Prior and PMF-GRN is motivated by their complementary roles in the workflow. GLM-Prior uses paired motif and gene-body sequence information to construct a sequence-derived scaffold of putative regulatory edges. This scaffold can be generated as a general prior matrix or tailored to specific cell lines or conditions when appropriate training data are available. However, as GLM-Prior has not seen any experimental data, its predictions require refinement using context-specific observations. PMF-GRN provides this refinement by integrating single-cell expression data within a principled generative framework, updating edge probabilities based on observed regulatory patterns while maintaining consistency with the sequence-derived prior. Moreover, like most GRN inference algorithms, PMF-GRN relies on prior knowledge to obtain a meaningful posterior GRN; without an informative prior, factor identifiability issues limit interpretability. GLM-Prior helps mitigate this limitation by providing TF-gene structural information, while PMF-GRN tailors the prior to the data, quantifies uncertainty, and produces context-specific networks. Together, this integration yields GRNs conditioned on both sequence-derived priors and single-cell expression data.

### GLM-Prior’s generalization scales with data composition across species and cell types

We assess GLM-Prior by benchmarking across six cell lines spanning yeast, human, and mouse (Figure 2). For each organism, we create gene-TF sequence pairs by extracting nucleotide sequences for genes from GTF annotations and TF binding motifs from CisBP (22). We then assign regulatory labels to these pairs using available interaction databases: YEASTRACT (11) for yeast, and STRING (23; 24; 25; 26) and TRRUST (27; 28) for human and mouse. These databases provide curated positive regulatory associations, but do not define an exhaustive set of true negative TF-gene pairs. We therefore treat gene-TF pairs without an annotated interaction as unlabeled negatives for training, recognizing that some sampled negatives may correspond to regulatory interactions that are absent from current databases or active only in specific cellular contexts.

**Figure 2:**
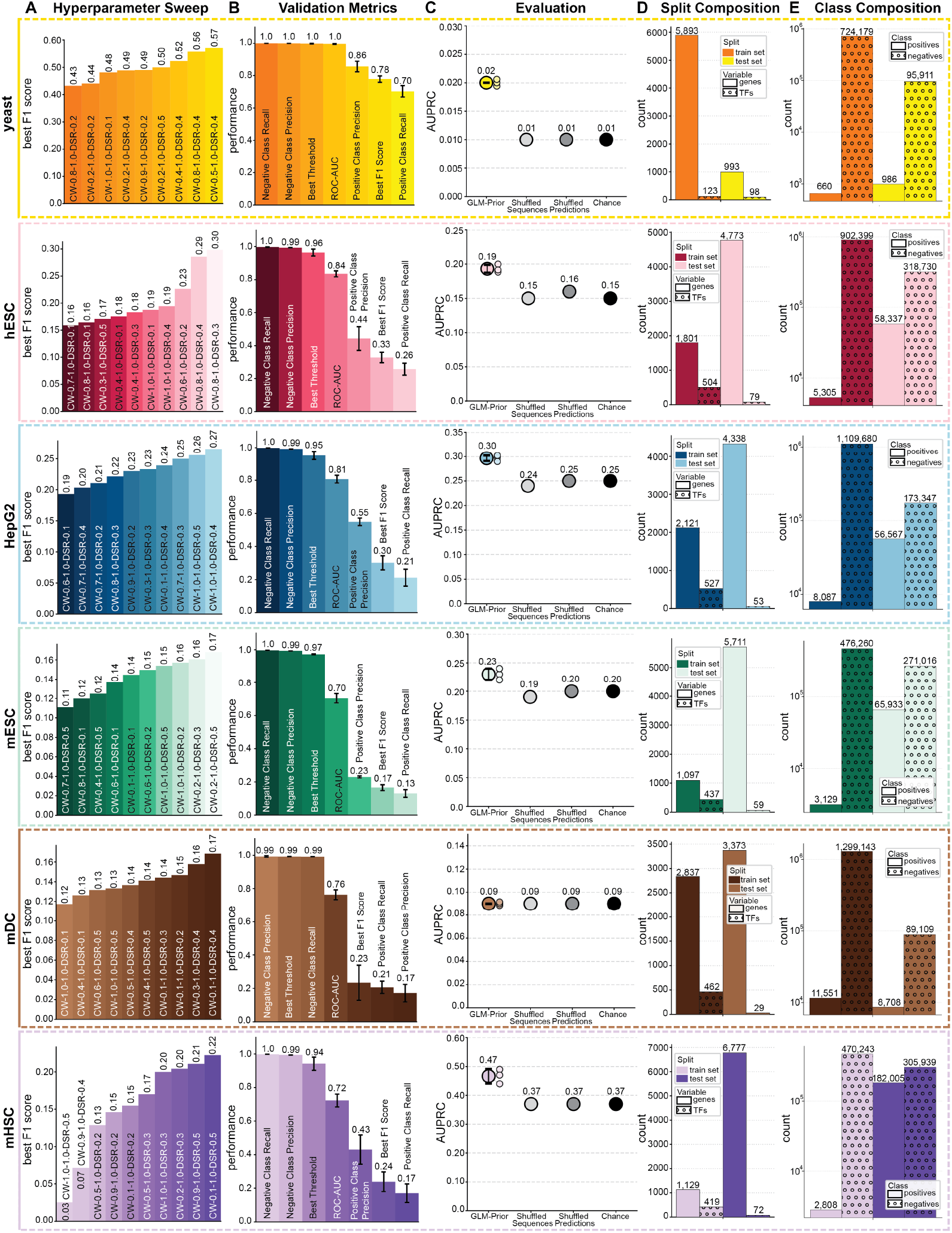
GLM-Prior model performance across yeast, hESC, HepG2, mESC, mDC, and mHSC cell lines. (**A**) One-epoch hyperparameter sweep over class weights and negative-class downsampling rates, evaluated by F1 score. (**B**) Validation metrics during 10 epoch training using the best hyperparameter configuration, trained across three random seeds (42, 43, and 44); bars summarize mean validation performance across seeds and error bars indicate ±1 standard deviation. (**C**) Test AUPRC on held-out reference networks with performance shown across three random seeds, with points indicating individual seed values and summary marks indicating mean performance with observed seed-to-seed variability. Baseline controls are shown as single reference points: light gray represents inference on shuffled sequences, dark gray represents shuffled predictions, and black represents chance performance based on the positive-label rate. (**D**) Train and test split composition, shown as the number of genes and TFs. (**E**) Class composition, shown as the number of positive and negative labels in train and test sets.

To assess out-of-distribution performance in this supervision setting, we evaluate on held-out test sets constructed from independent reference networks, including the literature-curated reference from (29) for yeast, and cell-line-specific BEELINE-derived ChIP-seq reference networks (30) for human and mouse. Before training each cell-line-specific model, we ensure strict independence between training and test sets at the level of gene-TF sequence pairs and their labels, eliminating sequence and label leakage. This design ensures that performance reflects the model’s ability to generalize to unseen sequences and correctly predict their interaction labels.

To evaluate model performance within each cell line, we implement a multi-stage benchmarking procedure (Figure 2A-E). In Figure 2A, we perform a one-epoch hyperparameter sweep over combinations of class weights in the loss function and downsampling rates for the negative class. Negative-class downsampling is necessary to make training computationally tractable given the combinatorial number of possible gene-TF pairs; however, this sampling strategy may exclude unlabeled pairs that correspond to unannotated regulatory interactions, preventing the model from learning from those examples during training. We identify the configuration that yields the highest F1-score and use these parameters for full model training. In Figure 2B, we train the model for ten epochs using this selected configuration and monitor performance on a heldout validation set. To assess sensitivity to model initialization, we repeat final training across three random seeds and report seed-level variability.

Following training, we perform inference separately for each seed-specific model across all gene-TF pairs, including pairs unseen during training, and calculate AUPRC for each resulting prior network against the corresponding held-out reference network labels (Figure 2C). In addition to the positive-rate chance baseline, we report two negative controls: (*i*) inference on test sets with gene and TF sequences shuffled to disrupt sequence-label correspondence while preserving class composition, and (*ii*) shuffled prediction scores on the original test set, which removes ranking signal and should approach the positive-rate baseline. To interpret what drives the resulting performance, we examine the number of genes and TFs represented in the training and test sets for each cell line (Figure 2D), providing context for possible domain shifts between splits. Finally, we quantify the class composition of positive and negative examples in the training and test sets (Figure 2E), allowing us to relate generalization behavior to the underlying data balance.

Across the six evaluated cell lines, GLM-Prior shows context-dependent agreement with held-out reference networks, with performance varying alongside the size and composition of the training and test data (Figure 2D-E). In yeast, GLM-Prior achieves near-chance AUPRC across all three random seeds (0.02 for each seed) compared to 0.01 for the baseline controls, despite strong within-training validation metrics (Figure 2B). This limited out-of-distribution agreement likely reflects the extreme class imbalance in the training data, where only 660 positive interactions are available among 724, 179 negatives. Although the held-out reference contains 986 positive labels, the small number of positive training examples and high validation performance suggest that the model may over-fit regulatory patterns present in the training data without transferring strongly to held-out gene-TF combinations.

In human embryonic stem cells (hESC), GLM-Prior shows a modest above-baseline signal, with AUPRC values ranging from 0.19 to 0.20 across three seeds, compared with 0.15 for chance and shuffled-sequence controls and 0.16 for shuffled predictions. This improvement is small but consistent across random initializations. hESC includes a larger pool of positive training examples (5, 305) than yeast, and the held-out reference network contains many labeled positives. However, TF coverage drops substantially from 504 TFs in training to 79 TFs in testing, indicating that performance is evaluated over a narrower set of TFs than those available during training. We therefore interpret hESC performance as modest evidence of agreement with the held-out reference network, rather than as strong evidence of broad mechanistic generalization.

In HepG2, GLM-Prior achieves higher AUPRC values than in hESC, ranging from 0.29 to 0.30 across seeds, compared with 0.25 for chance and shuffled predictions and 0.24 for shuffled sequences. This performance is supported by a larger number of positive training labels (8, 087) and a well-represented held-out reference set (56, 567 positives). Although the held-out set contains fewer TFs than the training set (53 versus 527), GLM-Prior consistently exceeds the negative controls. These results suggest that the model captures sequence-associated signal that improves ranking of labeled interactions in this cell line, while remaining limited to agreement with the available reference annotations.

In mouse embryonic stem cells (mESC), GLM-Prior achieves AUPRC values ranging from 0.22 to 0.24 across seeds, compared with 0.20 for chance and shuffled predictions and 0.19 for shuffled sequences. This represents a modest but reproducible improvement over the controls. The training set includes 3, 129 positives among 476, 260 negatives, while the held-out reference set is substantially more balanced (65, 933 positives and 271, 015 negatives) and covers 5, 711 genes unseen during training. These results indicate reproducible above-baseline agreement with the held-out reference network across unseen genes, with the modest margin over baseline suggesting that performance is measurable but limited in magnitude.

In mouse dendritic cells (mDC), GLM-Prior remains at chance across all three seeds, with AUPRC of 0.09 for GLM-Prior and all baseline controls. This indicates that the trained sequence model does not provide detectable ranking signal beyond the positive-label rate for this held-out reference. Although the mDC training set contains a large number of positive labels (11, 551), TF coverage drops sharply from 462 TFs in training to 29 TFs in testing. This difference may contribute to poor generalization, but the result primarily indicates that GLM-Prior does not improve agreement with the available mDC reference network under this evaluation design.

Finally, mouse hematopoietic stem cells (mHSC) show the strongest above-baseline performance, with GLM-Prior AUPRC values ranging from 0.44 to 0.49 across seeds, compared with 0.37 for chance, shuffled sequences, and shuffled predictions. The observed improvement is larger than in the other mammalian contexts, although seed-to-seed variation is also more apparent. The held-out reference set is large (182, 005 positives and 305, 939 negatives) and includes 6, 777 genes and 72 TFs, providing a broad evaluation set. These results indicate that GLM-Prior improves the ranking of labeled interactions in the held-out reference network, producing the strongest above-baseline performance among the evaluated cell lines.

Together, these results show that GLM-Prior’s performance depends strongly on the composition and coverage of the available training labels and held-out reference networks. Across seeds, the clearest improvements over negative controls occur in HepG2 and mHSC, while hESC and mESC show modest but reproducible gains, yeast remains close to chance, and mDC shows no detectable improvement. We therefore interpret above-chance performance as evidence that GLM-Prior can learn sequence-associated signals that improve agreement with held-out reference annotations in some contexts.

To evaluate whether this signal could instead be attributed to simpler properties of the input data, we performed diagnostic analyses of gene body composition, sequence similarity, gene length, and database annotation features (Appendix Section A.4). These diagnostics address distinct potential sources of proxy learning: gene body composition and length test whether predictions are dominated by broad sequence-scale properties, sequence similarity tests whether performance is explained by sequence relatedness among genes or gene-family structure, and annotation-based analyses test whether predictions are driven by uneven representation of genes or TFs in the interaction databases.

Across these analyses, GLM-Prior’s predictions are not driven solely by broad compositional differences among gene bodies, sequence relatedness among genes, gene length, or database annotation density. This strengthens the interpretation of the main results by showing that the above-baseline agreement with held-out reference networks cannot be reduced to these simpler properties of the input data.

### GLM-Prior supports mammalian transfer learning and multi-species prior construction

Following our six cell line evaluation of GLM-Prior, we next ask whether a sequence-derived model trained in one species can be used to construct prior knowledge in a species not seen during training (Figure 3). This is a central motivation for GLM-Prior: existing prior construction strategies often require species- or cell-line-specific experimental data, limiting their use in systems where regulatory annotations, chromatin accessibility profiles, or matched expression datasets are unavailable. A transferable sequence-based model could provide an initial putative TF-gene prior in such settings, which can then be evaluated or refined when reference networks or expression data become available.

**Figure 3:**
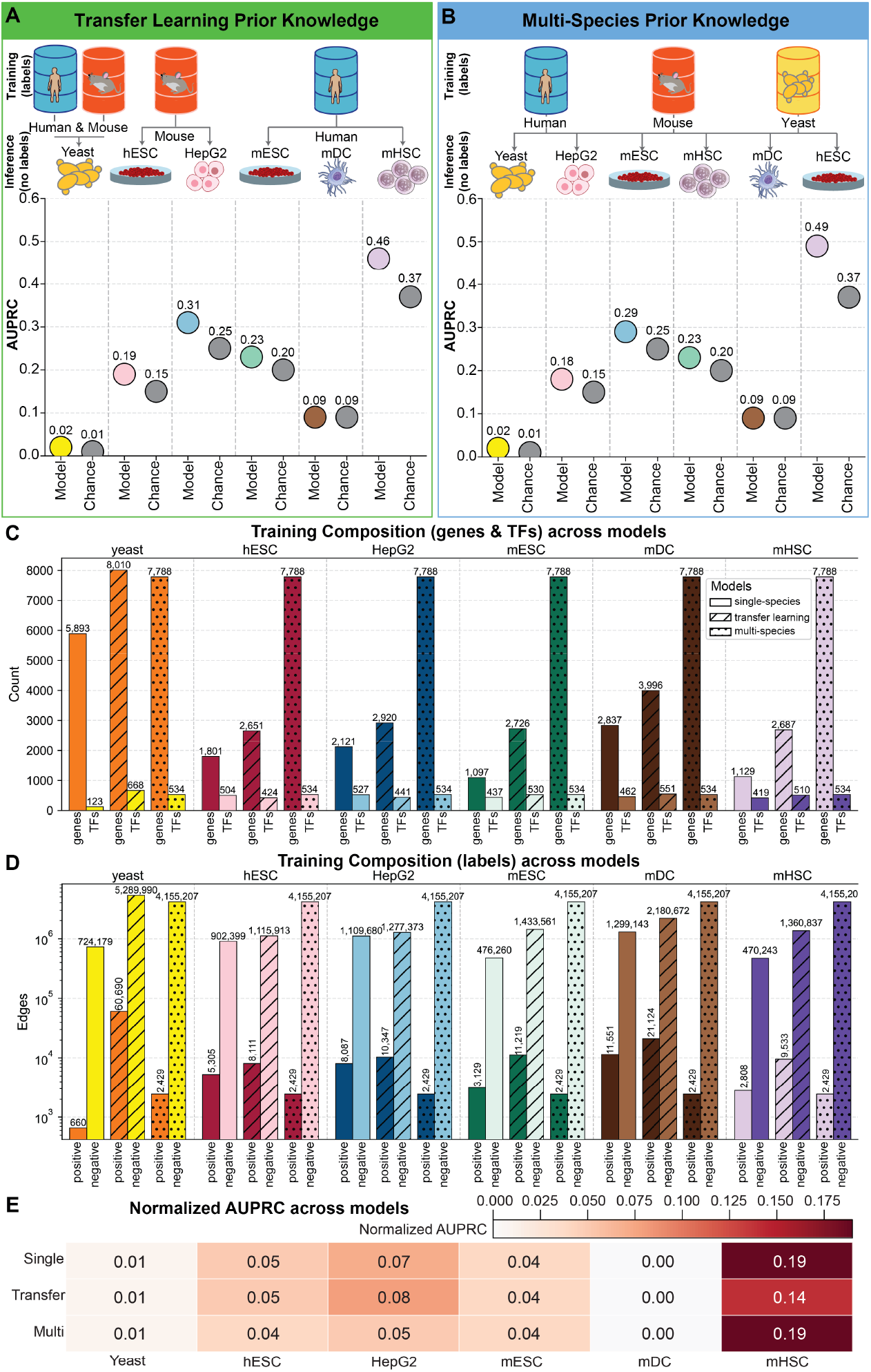
Transfer learning and multi-species training with GLM-Prior. (**A**) Schematic and AUPRC results of the species-transfer learning setup, showing three configurations: (*i*) a model trained jointly on human and mouse data evaluated on yeast, (*ii*) a mouse-only model evaluated on human cell lines, and (*iii*) a human-only model evaluated on mouse cell lines. (**B**) Schematic and AUPRC results of the multi-species model trained sequentially on human, mouse and yeast data, evaluated across six cell lines. (**C**) Comparison of training composition, shown as the number of genes and TFs, for single-species, species-transfer, and multi-species models. (**D**) Class composition, shown as the number of positive and negative labels, in the corresponding training sets. (**E**) Normalized AUPRC across single-species, species-transfer, and multi-species models for each cell line.

Building on the previous section, which established that generalization depends strongly on data composition, we test whether GLM-Prior preserves predictive signal across species with different degrees of evolutionary relatedness. We perform two complementary experiments: A) transfer learning, where models trained in one species are applied to another, and B) multi-species training, where a single model is trained sequentially on human, mouse, and yeast data. In all cases, held-out evaluation sets are fully disjoint from training data, and overlapping gene and TF identifiers are removed to prevent cross-species leakage.

In Figure 3A, we provide a schematic for the transfer learning setup. Three transfer learning configurations are evaluated, (*i*) a model trained jointly on human and mouse data, evaluated on yeast, (*ii*) a mouse-only model evaluated on human cell lines (hESC and HepG2), and (*iii*) a human-only model evaluated on mouse cell lines (mESC, mDC, and mHSC). Across all six cell lines, we find that transfer learning maintains or modestly improves performance relative to single-species models.

In yeast, performance remains near chance (AUPRC = 0.02 vs 0.01), indicating that transfer from mammalian training data does not provide a useful prior for this more distantly related organism under the current evaluation. In contrast, transfer between human and mouse is more encouraging, where the mouse-trained model maintains performance in hESC (AUPRC = 0.19) and improves performance in HepG2 (AUPRC = 0.31), while human-trained models applied to mouse cell lines achieve performance comparable to the corresponding single-species models. These results suggest that GLM-Prior can construct informative sequence-derived priors across closely related mammalian species, while also showing that transfer does not extend uniformly across more evolutionarily distant systems.

In Figure 3B, we present the multi-species model trained sequentially on human, mouse, and yeast data and evaluated on all six cell lines. The multi-species model achieves comparable AUPRCs relative to both single-species and species-transfer settings (yeast = 0.02, hESC = 0.19, HepG2 = 0.29, mESC = 0.23, mDC = 0.09, and mHSC = 0.49).

To interpret these trends, Figure 3C-D summarize the training compositions and normalized AUPRC (Figure 3E) across single, transfer, and multi-species setups. Across each cell line, transfer learning consistently expands the number of genes, TFs, and positive labels seen during training. In some settings, such as with HepG2, this broader coverage allows for a modest normalized AUPRC increase (0.08 vs 0.07 single-species). In others, transfer learning maintains single-species performance. In contrast, mHSC shows a slight decrease in performance (0.14 species-transfer vs 0.19 single-species) despite larger training breadth. This drop suggests that the regulatory logic active in mouse HSCs is relatively cell-type and species-specific and therefore not well represented in the context-agnostic human interaction data used for species-transfer training, limiting the model’s ability to fully recover mHSC-specific edges.

Notably, the multi-species model maintains similar performance to the single-species models, with only slight performance drops in hESC and HepG2. This result suggests that multi-species training can produce a broadly applicable sequence-derived prior without requiring a separate model for each target context. Such a model may be useful when the target species or cell type is poorly characterized, although its utility remains dependent on the evolutionary and regulatory similarity between the training data and the target system.

Together, these results show that GLM-Prior provides a practical route for constructing prior knowledge in settings where species- or cell-line-specific regulatory data are limited. Transfer between human and mouse maintains or modestly improves agreement with held-out reference networks, supporting the use of GLM-Prior for mammalian transfer-learning scenarios. However, transfer to yeast remains near chance, indicating that this capability is context-dependent and does not establish universal transfer across distant species. Multi-species training further suggests that a single sequence-derived model can provide stable priors across the evaluated contexts, while highlighting the need for additional reference networks to assess transferability across a broader range of organisms and cell types.

### Integrating GLM-Prior into PMF-GRN supports prior-conditioned GRN refinement

To evaluate how sequence-derived prior knowledge interacts with expression-based GRN inference, we next used each cell line-specific GLM-Prior matrix as input to PMF-GRN. Whereas Stage 1 (Section 1) scores TF-gene pairs from sequence to construct a prior matrix, Stage 2 (Section 1) treats these predictions as structural prior knowledge and refines the network using single-cell expression data. For each cell line, we paired GLM-Prior with matched single cell datasets, including GSE125162 (31) and GSE144820 (32) for yeast, and the corresponding expression datasets provided in BEELINE (30) for the human and mouse cell lines. This experiment assesses whether expression-based inference improves agreement with held-out reference networks, and how much PMF-GRN modifies the sequence-derived prior during inference.

Across all datasets, integrating GLM-Prior into PMF-GRN results in equal or improved AUPRC relative to the prior alone (Figure 4A). The magnitude of improvement varies by species and cell line, where performance remains unchanged in yeast (0.02) and hESC (0.19), while gains are observed in HepG2 (0.30 prior → 0.38 GRN), mESC (0.23 prior → 0.27 GRN), and mHSC (0.49 prior → 0.52 GRN). These changes occur despite PMF-GRN modifying only a small fraction of the GLM-Prior edge matrix (Figure 4B), with added edges ranging from 0.03% in yeast to 0.54% in mHSC, and edge removals remaining minimal across all contexts. The modifications are biased toward edge additions rather than deletions, suggesting that PMF-GRN primarily preserves the GLM-Prior scaffold while adding a limited number of expression-supported edges.

**Figure 4:**
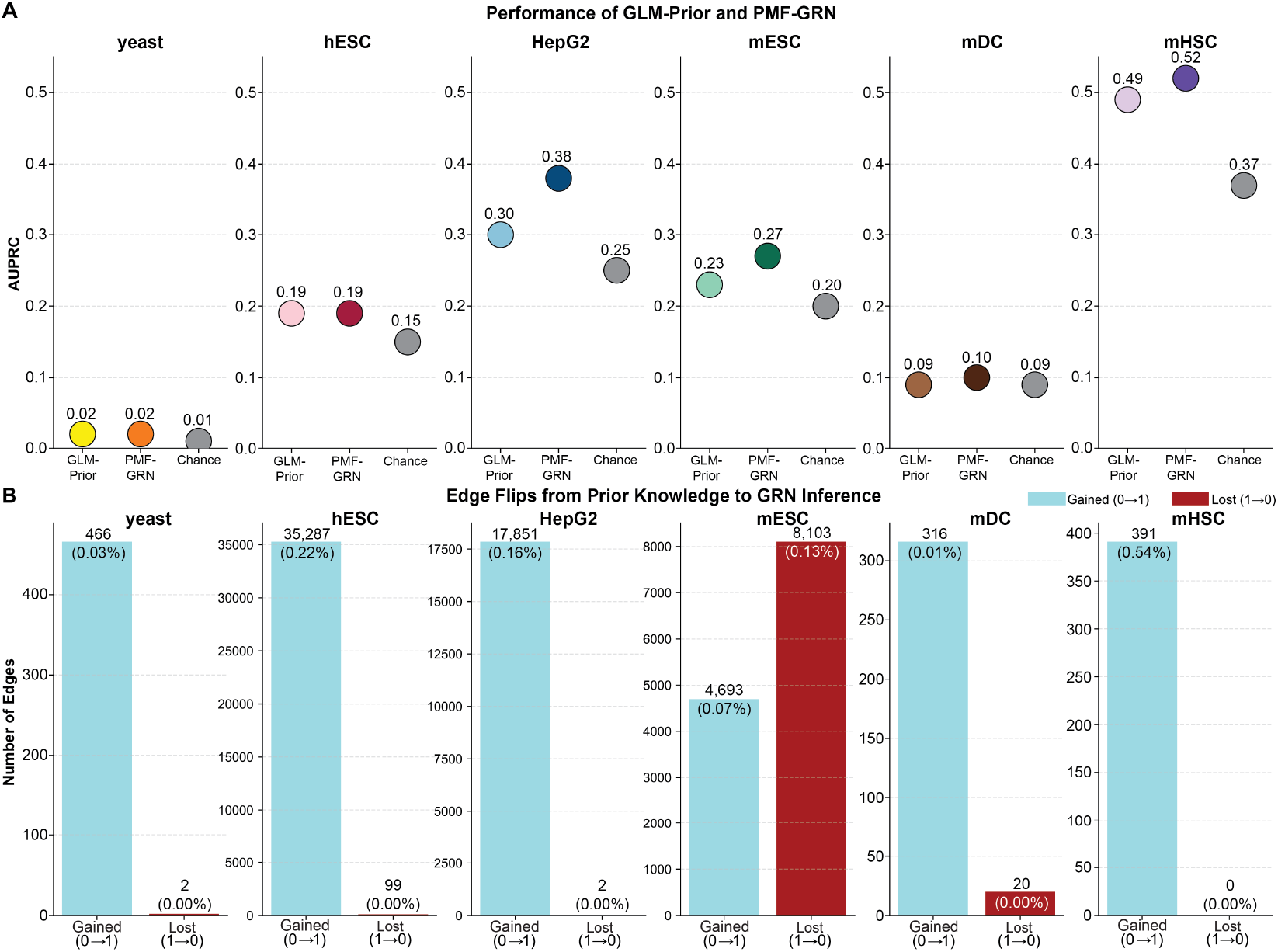
Integration of GLM-Prior with PMF-GRN for downstream GRN inference. (**A**) AUPRC comparison of GLM-Prior alone versus integrated with PMF-GRN across six cell line contexts. (**B**) Edge flip analysis quantifying changes made to the prior knowledge matrix during PMF-GRN inference. Blue bars indicate prior edges added and red bars indicate prior edges removed during inference.

The extent of structural change does not strictly track with performance improvement. For example, hESC and mESC exhibit tens of thousands of added or removed edges, yet show minimal differences in AUPRC. This mismatch suggests that many of the changes introduced by PMF-GRN occur outside the held-out evaluation portion of the network, which covers only a subset of TF-gene interactions. Conversely, in yeast, where GLM-Prior is close to chance (AUPRC = 0.02) and the available training interactions are sparse, PMF-GRN does not improve agreement with the held-out reference network despite access to two large single-cell expression datasets. This result suggests that expression-driven inference alone cannot compensate when the input prior has limited alignment with the available reference annotations.

These findings indicate that expression-based inference provides prior-dependent refinement rather than a wholesale reconstruction of the network. When the sequence-derived prior demonstrates above-baseline agreement with the held-out reference network, PMF-GRN generally preserves most prior edges and introduces limited expression-supported modifications. In this dual-stage framework, the quality of the GLM-Prior matrix strongly influences the achievable reference-network agreement, while PMF-GRN contributes selective improvements where transcriptomic variation provides additional support. When the prior is near chance, as in yeast, expression-based GRN inference has limited capacity to recover agreement with the available reference network.

### Comparing prior construction strategies across yeast, mouse, and human cell lines

Next, we assess how GLM-Prior compares to widely used accessibility-based prior construction approaches, evaluating performance relative to CellOracle and Inferelator priors across six cell line contexts. Unlike GLM-Prior, which uses sequence-level information to score TF-gene pairs, both CellOracle and Inferelator derive putative interactions from chromatin accessibility and motif enrichment. Importantly, these accessibility-based priors are constructed using ATAC-seq measured in the matched cell line, whereas GLM-Prior is trained on context-agnostic interaction resources (STRING and TRRUST) and evaluated without explicitly conditioning on cell-line specific chromatin state. This comparison therefore evaluates whether sequence-derived priors can match or exceed accessibility-based priors in agreement with available reference networks across diverse biological settings.

Across all six cell lines, GLM-Prior performs at or above chance, exceeding chance performance in five of the six contexts(Figure 5A). In comparison, Inferelator-Prior performs above chance in two out of six cell lines, and CellOracle’s Prior achieves above-chance performance in three out of six, indicating that prior performance varies substantially across methods and contexts. Yeast represents the main exception, where CellOracle produces the highest-performing prior (AUPRC 0.05 versus 0.03 for Inferelator-Prior and 0.02 for GLM-Prior). This result is consistent with the structure of accessibility-based priors, which are well suited to promoter-proximal, motif-supported regulatory assignments in yeast. In this setting, GLM-Prior is further limited by the small number of positive regulatory labels (660) available for model training (as seen in Figure 2E).

**Figure 5:**
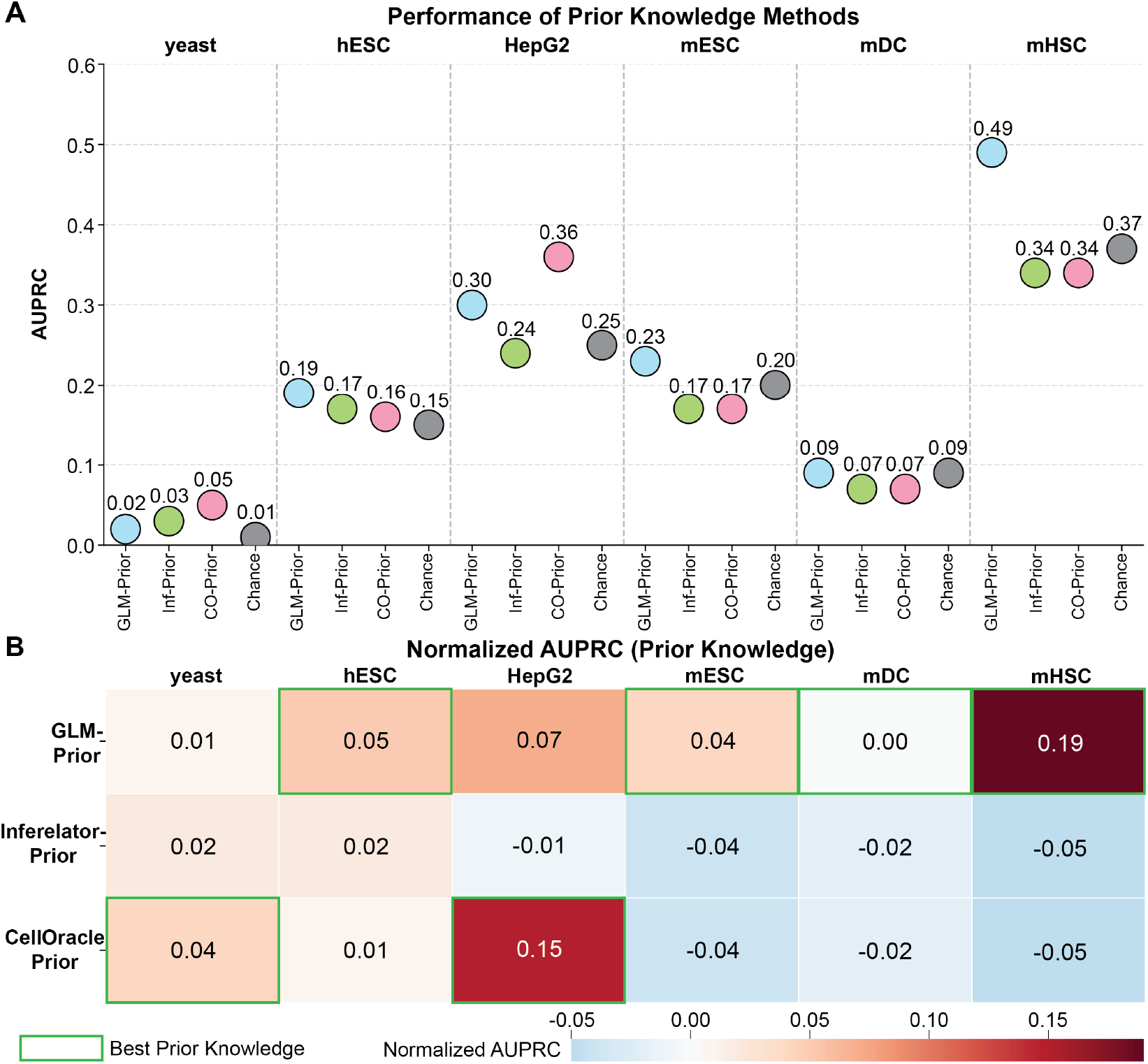
Comparison of prior construction strategies across species. (**A**) AUPRC of GLM-Prior, Inferelator-Prior and CellOracle Prior across six cell lines, with chance performance (gray) provided for each context. (**B**) Normalized AUPRC values with the best-performing prior (green box) highlighted in each cell line.

A broader comparison across species using normalized AUPRC values further illustrates these patterns (5B), where normalized AUPRC is defined as (AUPRC − chance)*/*(1 − chance). While GLM-Prior provides the most consistent gains overall, CellOracle’s prior outperforms all other approaches in HepG2 (0.36 versus 0.30 for GLM-Prior and 0.24 for Inferelator-Prior). This comparatively strong performance in a mammalian cell line may reflect alignment between the construction of CellOracle’s prior and the BEELINE reference network used for evaluation. BEELINE assembles reference networks using ChIP-seq supported TF-gene edges curated from ENCODE (10), ChIP-ATLAS (33) and ESCAPE (34) databases, and HepG2 is among the most densely profiled ENCODE cell lines. As a result, the HepG2 reference likely contains a large and internally consistent set of TF occupancy-supported edges that are more recoverable by accessibility-conditioned, proximity-based linking, which is the type of evidence CellOracle prioritize when constructing its prior from motifs in accessible regions. By contrast, Inferelator-Prior follows a related accessibility- and motif-based strategy but imposes stronger sparsity which may remove a substantial number of true TF-gene pairs represented in the HepG2 reference network and thereby reduce its alignment with that reference.

In contrast, GLM-Prior is the top-performing method in the four remaining cell lines: hESC, mESC, mDC, and mHSC. In hESC, all methods exceed chance, with GLM-Prior achieving the highest AUPRC (0.19 versus 0.17 for Inferelator-Prior and 0.16 for CellOracle’s prior). In mESC and mHSC, the advantage of GLM-Prior becomes more pronounced, as accessibility-based priors fall below chance while GLM-Prior remains predictive. These results suggest that sequence-derived priors can provide stronger agreement with held-out reference networks than accessibility-based priors in several mammalian contexts, despite not using matched cell-line-specific chromatin accessibility data.

These comparisons show that prior construction strategies have context-dependent performance. Accessibility-driven approaches such as CellOracle can perform well when the reference network aligns with motif- and accessibility-based edge construction, as observed in yeast and HepG2. GLM-Prior, by contrast, provides a sequencing technology-independent alternative that achieves the strongest agreement with held-out reference networks in four of five mammalian contexts. The complementary performance profiles across species and cell lines indicate that no single prior construction approach is universally optimal, but that sequence-derived priors can provide a scalable strategy for prior construction when matched chromatin accessibility data are unavailable or when transfer across related mammalian contexts is desired.

### Paired prior-GRN strategies reveal prior-dependent gains from expression-based inference

Following our comparative analysis of prior construction approaches, we next asked how each complete prior-GRN pipeline performs across the six cell lines (Figure 6). GLM-Prior and PMF-GRN form our dual-stage pipeline, combining a finetuned sequence language model for prior construction with a probabilistic matrix factorization model for GRN inference from single cell expression. CellOracle constructs a prior (“CellOracle baseGRN”) using motif enrichment within accessible promoter regions and pairs it with a Bayesian ridge regression model for GRN inference. Similarly, Inferelator-Prior uses chromatin accessibility and TF motif enrichment to build a prior edge matrix that is then refined using regularized regression during GRN inference. By comparing these three paired pipelines, GLM-Prior + PMF-GRN, Inferelator-Prior + Inferelator, and CellOracle prior + CellOracle, we ask two related questions: (*i*) how well each method converts its own prior into a GRN that agrees with the corresponding reference network, and (*ii*) to what extent expression-based GRN inference can compensate for weak priors versus refining already strong ones.

**Figure 6:**
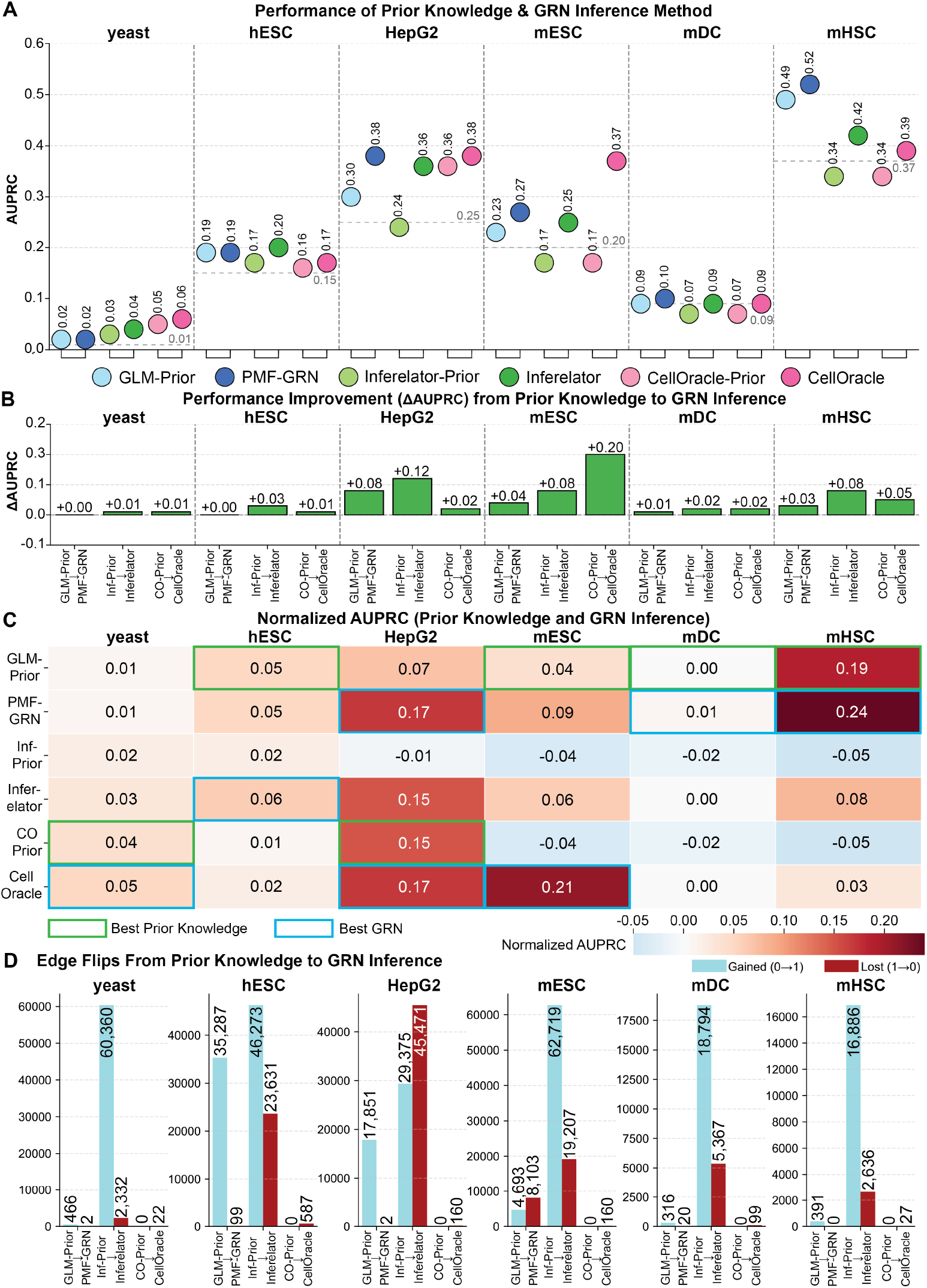
Paired prior-GRN strategies reveal prior-dependent gains from expression-based GRN inference across six cell lines. (**A**) AUPRC of each prior (GLM-Prior, Inferelator-Prior, and CellOracle’s prior) and its corresponding GRN (PMF-GRN, Inferelator, CellOracle) across six cell lines, with gray dashed lines indicating the chance level in each species. (**B**) Change in AUPRC from prior to GRN (ΔAUPRC) for each paired method, quantifying the benchmark gain from expression-based inference. (**C**) Normalized AUPRC performance for priors and GRNs in each cell line. Green boxes highlight the best prior and blue boxes highlight the best GRN per cell line. (**D**) Edge flips between prior and GRN for each method, showing how GRN inference methods modify their priors.

To interpret these patterns, we consider the absolute AUPRCs in Figure 6A together with the corresponding ΔAUPRC values in Figure 6B. In yeast and mDC, all priors are at or only slightly above chance, and GRN inference yields minimal gains (ΔAUPRC ≤ 0.02), indicating that expression-based models show limited ability to improve reference-network agreement when the starting prior is near-random. In hESC and mHSC, GLM-Prior already achieves above-chance performance, and PMF-GRN adds only small improvements, whereas Inferelator and CellOracle modestly refine their weaker priors. These contexts illustrate that when priors are already above baseline, GRN inference primarily fine-tunes edge weights rather than substantially reshaping the network.

By contrast, HepG2 and mESC are the settings where GRN inference provides the largest benefits. Here, priors span a range from near- or slightly below-chance (Inferelator-Prior in HepG2; Inferelator-Prior and CellOracle prior in mESC) to moderately above-chance (GLM-Prior and CellOracle prior in HepG2; GLM-Prior in mESC), leaving partially specified structure that expression data can refine. In HepG2, GLM-Prior + PMF-GRN and Inferelator-Prior + Inferelator show sizeable increases in AUPRC, while CellOracle begins from the strongest prior and improves only modestly. In mESC, regression-based inference on top of weak or near-chance priors yields the single largest observed ΔAUPRC for CellOracle and substantial gains for Inferelator, whereas PMF-GRN makes only a small but consistent improvement over an already above-chance GLM-Prior. Taken together, these six cell lines reveal a coherent pattern: the magnitude of GRN-inference gains is strongly prior-dependent, with the largest AUPRC gains arising when priors are of intermediate quality, limited performance improvements when priors are very weak, and only modest refinements when priors are already strong.

We next normalize AUPRC by the chance level in each species (as in Section 1) to directly compare performance above (or below) chance across methods and cell lines (Figure 6C). In this view, GLM-Prior provides the best prior in four of six cell lines (hESC, mESC, mDC, mHSC), while CellOracle prior yields the strongest priors in yeast and HepG2. We observe that in four mammalian cell lines (HepG2, mESC, mDC, and mHSC), Inferelator-Prior rarely exceeds random-chance performance and typically remains close to or below chance. Similarly, in all three mouse cell lines, CellOracle’s prior does not construct a prior above chance. These patterns suggest that proximity- and accessibility-based priors do not consistently align with the held-out mammalian reference networks used here, particularly in the mouse cell lines.

In addition, we find that the best GRNs largely track the best priors, with PMF-GRN producing the top-performing GRN in three of six cell lines, CellOracle in two, and Inferelator in one (Figure 6C). Notably, there are instances where GRN inference can partially compensate for a weaker prior. In mESC, CellOracle’s prior is below chance (normalized AUPRC = −0.04), yet its regression-based GRN achieves the highest normalized performance across all methods (0.21), outperforming GLM-Prior + PMF-GRN despite starting from a lower-performing prior. This behavior is clarified by examining how each GRN model modulates its prior in terms of edge flips (Figure 6D).

PMF-GRN primarily improves performance by adding edges to the prior, expanding the inferred network while leaving most prior edges unchanged. Inferelator both adds and removes edges, suggesting a balance between expansion and pruning. In contrast, CellOracle’s regression-based design only allows removal of edges: it can shrink edge weights to zero but cannot introduce edges that were absent from the accessibility-derived prior. The large performance gains observed for CellOracle in settings such as mESC therefore arise from pruning a dense prior rather than adding new edges during GRN inference. This pruning behavior can make a low-performing but dense prior align more strongly with the held-out reference network after GRN inference, yet also highlights an important limitation: when single-cell expression is used only to filter a fixed network, not to propose new edges, it cannot expand the regulatory scaffold beyond what is already encoded in the prior.

Together, these results show that while GRN inference can substantially improve performance in some settings, particularly when priors are neither near-random nor already highly aligned with the reference network, the overall ranking of methods is largely dictated by prior quality. Expression-based GRN inference models mostly refine and re-weight the structure supplied by the prior, rather than overturning it, underscoring the central role of prior construction in determining benchmarked GRN performance.

### Cross-method comparison disentangles prior and GRN inference contributions to performance

To disentangle the contribution of prior construction and GRN inference to reference-network agreement, we next performed a full cross-comparison in which each GRN inference method is applied to each prior across all six cell lines (Figure 7). Specifically, we combine GLM-Prior, Inferelator-Prior, and CellOracle baseGRN with PMF-GRN, Inferelator, and CellOracle, yielding nine prior-GRN combinations per cell line. In Figure 7A, we show the resulting AUPRC values, along with species-specific chance levels (gray dashed line), and the baseline performance of the prior (colored dashed line) used for GRN inference. In Figure 7B, we report the same results normalized relative to chance, where values above zero indicate performance above random. We focus our interpretation on the normalized AUPRC values, which allows for easier comparison across species and methods, and refer to the raw AUPRCs in Figure 7A where relevant.

**Figure 7:**
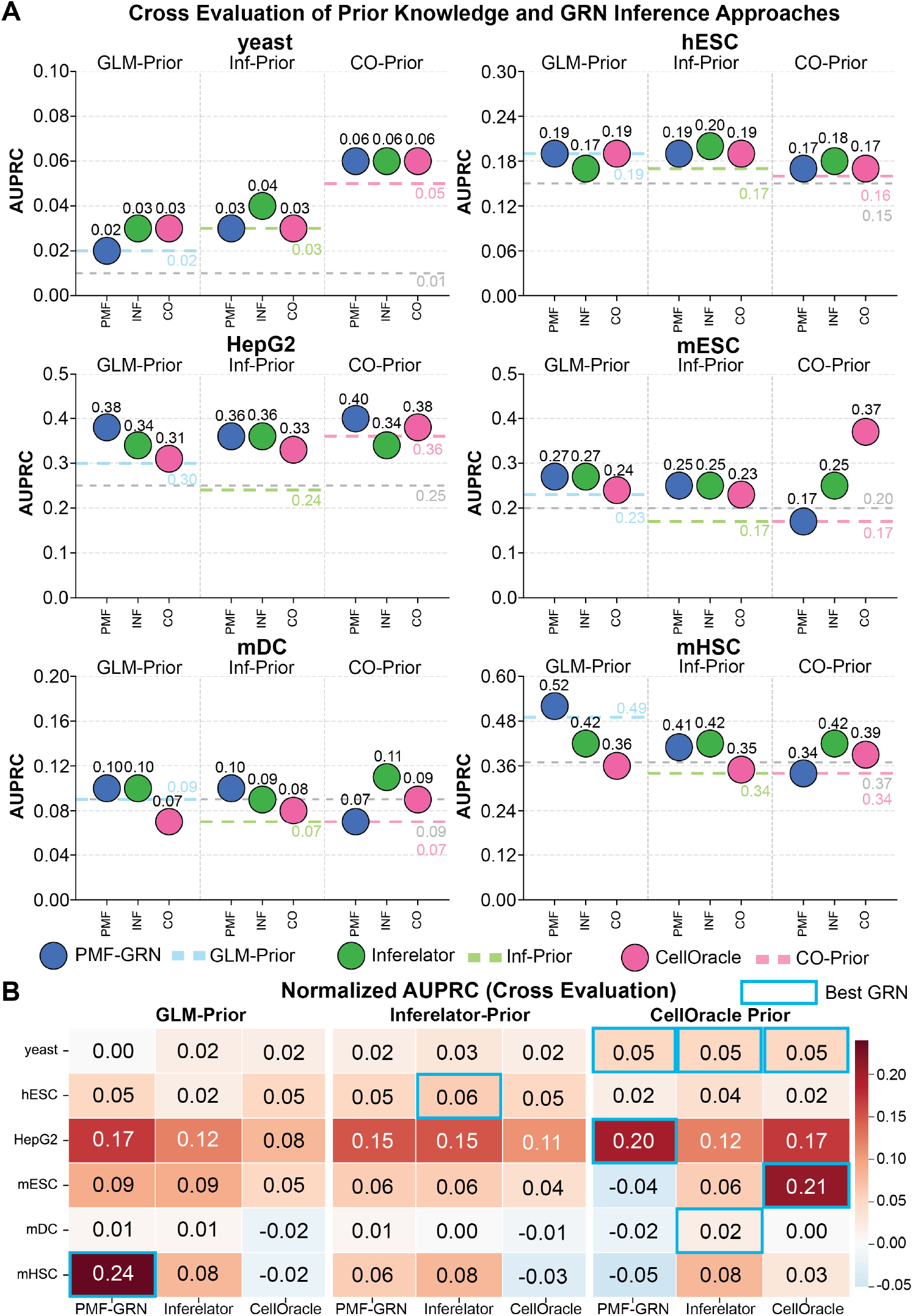
Cross-method comparison disentangles prior and GRN inference contributions. **(A)** AUPRC for all nine combinations of prior and GRN inference methods across six cell lines. Gray dashed lines indicate chance performance, while light blue, light green, and light pink dashed lines represent baseline performance of GLM-Prior, Inferelator-Prior, and CellOracle prior, respectively, across each cell line. **(B)** Normalized AUPRC of all nine combinations of prior and GRN for directly comparing across method and cell line. Blue boxes indicate highest GRN performance per cell line.

Across species, the dominant pattern is that the highest-performing inferred GRN for a given cell line is largely determined by the underlying prior, not by the choice of GRN inference method. In yeast, for example, any GRN method paired with CellOracle’s prior attains the same normalized AUPRC (0.05), whereas all combinations using GLM-Prior or Inferelator-Prior remain at or below 0.03. Once CellOracle’s prior is selected, the choice among PMF-GRN, Inferelator, or CellOracle has no quantifiable impact, indicating that the prior itself, rather than the choice of downstream inference algorithm, is the main constraint on benchmark performance.

Similar patterns appear in the mammalian cell lines. In hESC, GLM-Prior and Inferelator-Prior yield the highest-scoring priors (0.19 and 0.17 AUPRC, respectively), and nearly all GRNs inferred from these priors achieve competitive reference-network agreement. In HepG2, CellOracle’s prior is best (AUPRC = 0.36), and the two highest performing GRNs are obtained by pairing this prior with PMF-GRN or CellOracle, demonstrating again that performance is largely determined by the prior rather than the choice of inference algorithm. An analogous effect is observed in mHSC, where GLM-Prior combined with PMF-GRN yields the highest performing combination (AUPRC = 0.24), while all GRNs built on Inferelator-Prior or CellOracle’s prior plateau at 0.08 or lower. These results support a consistent conclusion: the quality and structure of the prior largely constrain achievable benchmark performance, while GRN inference methods primarily modulate performance within a range set by the prior.

The full cross-comparison design also allows us to ask whether any GRN inference method is inherently superior across priors and species. Across PMF-GRN, Inferelator, and CellOracle, we find no evidence that any single GRN inference method is uniformly superior. Each approach wins within different regimes defined by the combination of species and prior. PMF-GRN delivers the highest normalized performance in HepG2 when paired with CellOracle’s prior (0.20) and in mHSC when paired with GLM-Prior (0.24). PMF-GRN also performs well in yeast when paired with CellOracle’s prior, and mESC when paired with GLM-Prior or Inferelator-Prior. The Inferelator is best in hESC when paired with Inferelator-Prior (0.06) and in mDC when paired with CellOracle’s prior (0.02), and remains competitive in HepG2 across multiple priors. CellOracle is uniquely optimal in mESC when paired with CellOracle’s prior (0.21), substantially outperforming all other prior-GRN combinations in that species. The ranking of each GRN inference method thus changes with both prior and cell line, consistent with the interpretation that GRN methods act as context-dependent refiners of prior information using expression data, rather than serving as uniformly superior drivers of performance across settings.

The cross-method comparison further reveals characteristic interactions between each GRN inference method and the priors. PMF-GRN appears highly prior sensitive, where switching priors can produce large swings in performance within a given species. In mHSC, for instance, normalized performance for PMF-GRN ranges from −0.05 with CellOracle-Prior to 0.24 with GLM-Prior. In mESC, performance ranges from −0.04 with CellOracle Prior to 0.09 with GLM-Prior. This pattern indicates that PMF-GRN is most effective when initialized with priors that already show strong reference-network agreement, but has limited ability to improve priors that are near or below chance. The Inferelator, by contrast, is comparatively robust to prior choice. Within each species, the variation in normalized AUPRC across priors is typically modest (often within the range of 0.02 to 0.04). In mHSC, all three priors yield identical normalized performance (0.08) with the Inferelator. This suggests that the Inferelator relies more heavily on expression-based regularized regression, making it less sensitive to exact prior structure, while also limiting its ability to reach the highest benchmark performance in most cell lines. CellOracle exhibits strong method-prior compatibility in specific settings. For example, in mESC, CellOracle’s prior + Cell Oracle perform well (0.21), significantly outperforming all other combinations, indicating that the regression-based pruning used by CellOracle is particularly well matched to dense priors that require substantial edge removal to improve agreement with the reference network.

As each GRN inference method is applied to every prior, we can also ask whether cross-pairing ever allows a high-performing GRN inference method to elevate a lower-performing prior above combinations that use a higher-performing prior with a different inference method. Such reversals are rare. In mHSC, for example, no method applied to Inferelator-Prior (0) or CellOracle prior achieves performance comparable to GLM-Prior + PMF-GRN. In HepG2, there is a modest example of beneficial cross-pairing, where CellOracle prior + PMF-GRN achieves an AUPRC of 0.20, outperforming the original CellOracle prior + CellOracle pairing (0.17). The performance gains here are likely due to PMF-GRN adding expression-supported edges to the CellOracle prior. In mDC, CellOracle prior + Inferelator (0.02) is better than CellOracle prior + CellOracle or CellOracle prior + PMF-GRN (0.02), although all values remain near chance. These exceptions are incremental, not complete reversals of the prior-driven ranking, and reinforce the view that GRN inference can fine-tune benchmark performance but typically does not elevate a low-performing prior to the level of the highest-performing priors.

Overall, the cross-method comparison in Figure 7 supports three main conclusions. First, prior quality remains the primary determinant of benchmarked GRN performance, even when each GRN method is given access to each prior. Second, there is no universally better GRN inference algorithm: PMF-GRN, the Inferelator, and CellOracle perform best in different prior and species regimes, reflecting differences in how they use expression data to refine or re-weight the prior. Third, GRN inference methods modulate, rather than overturn, the structure imposed by the prior. Changing the inference algorithm can yield meaningful gains in some contexts, particularly when paired with an appropriate prior, but rarely elevates a weak prior to the level of a strong one. These findings reinforce the conclusion that prior construction is the main driver of reference-network agreement in this benchmark, with expression-based GRN inference primarily serving a prior-dependent refining role.

## Discussion

Constructing useful GRN models remains a central challenge in genomics, particularly in systems where regulatory annotations and matched experimental assays are incomplete or unavailable. In this work, we present GLM-Prior as a sequence-derived workflow for constructing TF-gene prior knowledge matrices and evaluate how these priors interact with downstream expression-based GRN inference. By integrating GLM-Prior with PMF-GRN, a probabilistic matrix factorization framework for single-cell expression data, we decouple prior construction from GRN inference and systematically assess how each component contributes to benchmarked GRN performance across yeast, mouse, and human cell-line contexts.

Our results first demonstrate that GLM-Prior performance is tightly linked to the composition of the available training data. Across six evaluated contexts, agreement with held-out reference networks scales with positive-label abundance, TF coverage, and the relationship between training and evaluation labels. Human and mouse cell lines with richer regulatory annotations support stronger above-baseline performance than yeast, where sparse positive labels and extreme class imbalance limit generalization despite strong within-training validation metrics. Here, hESC and mESC show modest but reproducible gains, HepG2 and mHSC show clearer improvements, and mDC remains at chance. These findings indicate that genomic language models can learn sequence-associated signals that improve agreement with available regulatory references in some contexts, but that their performance remains constrained by incomplete supervision and the coverage of current benchmark networks.

Transfer-learning experiments further support GLM-Prior as a workflow for constructing priors in settings where species- or cell-line-specific regulatory data are limited. Transfer between human and mouse maintains or modestly improves agreement with held-out reference networks compared to single-species models, suggesting that GLM-Prior can preserve useful sequence-associated predictive signal across related mammalian species. In contrast, transfer from mammalian training data to yeast remains near chance, indicating that this capability is context-dependent and does not extend uniformly across evolutionarily distant organisms. Further, multi-species training produces performance comparable to single-species or transfer models across the evaluated contexts, suggesting that a single sequence-derived model can provide stable priors without requiring separate model training for every target species. These results support the practical use of GLM-Prior for mammalian transfer-learning scenarios, while emphasizing that broader claims about transferability require additional reference networks across more organisms, cell types, and regulatory regimes.

Furthermore, benchmarking GLM-Prior against accessibility-based priors further shows that prior construction strategies have context-dependent strengths. Accessibility-driven approaches such as CellOracle perform well when their construction assumptions align with the reference network, as observed in yeast and HepG2. GLM-Prior, by contrast, achieves the strongest agreement with held-out reference networks in four of five mammalian contexts, despite not using matched cell-line-specific chromatin accessibility data. This comparison does not establish that sequence-derived priors recover specific classes of regulatory mechanisms, such as distal enhancer regulation or long-range TF-gene interactions. Rather, it shows that sequence-derived priors can provide competitive or stronger reference-network agreement in several mammalian settings and offer a scalable alternative when matched chromatin accessibility data are unavailable.

Integrating priors with multiple GRN inference algorithms reveals a consistent pattern: the quality and structure of the prior largely constrain achievable benchmark performance, while expression-based inference provides prior-dependent refinement. When GLM-Prior is used as input to PMF-GRN, AUPRC is equal to or higher than the prior alone across the evaluated contexts, with gains varying by cell line. Edge-flip analyses show that PMF-GRN generally preserves most prior edges and introduces limited expression-supported modifications, rather than reconstructing the network independently from expression data. The fully crossed comparison among GLM-Prior, Inferelator-Prior, CellOracle’s baseGRN, PMF-GRN, Inferelator, and CellOracle further shows that no GRN inference algorithm is uniformly superior across priors or species. Instead, the highest-performing inferred GRNs usually arise from combinations that use the highest-performing prior for that context. These results support the conclusion that prior construction is a major determinant of reference-network agreement in current GRN benchmarks, with downstream inference methods refining or reweighting the structure supplied by the prior.

Several limitations of our study point to important directions for future work. First, regulatory interaction databases are incomplete and noisy. Positive labels are derived from curated interaction resources, while pairs without annotated interactions are treated as unlabeled negatives for training. Some of these unlabeled negatives may correspond to real regulatory interactions that are absent from current databases or active only in specific cellular contexts. The use of negative-class downsampling in our GLM-Prior approach is necessary to make training computationally tractable given the combinatorial number of possible TF-gene pairs, but this sampling strategy may also exclude informative unlabeled pairs from training. Future work should explore sampling strategies and positive-unlabeled learning frameworks that better reflect the incompleteness of regulatory labels.

Second, evaluation is limited by the availability of benchmarkable reference networks. The six cell-line contexts used here represent the settings in which matched sequence inputs, training labels, and independent reference networks were available for systematic evaluation. These references provide useful standardized benchmarks, but they are not exhaustive maps of regulatory interactions. Expanding the number and diversity of reference networks will be essential for evaluating whether sequence-derived prior construction generalizes beyond the mammalian contexts where GLM-Prior performs best.

Third, the gene body input design remains an important area for further investigation. Gene body sequence provides a consistent anchor for constructing gene-level inputs, but it may also encode proxy features such as sequence composition, gene length, gene-family structure, or annotation density. To address this concern, we performed diagnostic analyses showing that GLM-Prior performance is not reducible to broad compositional differences among gene bodies, sequence relatedness among genes, gene length, or database annotation density. These analyses support the interpretation that GLM-Prior captures sequence-associated information beyond these simple proxy features, while leaving open the need for future model designs that incorporate richer regulatory context, including promoters, enhancers, chromatin state, three-dimensional genome organization, and cell-type-specific regulatory annotations.

Finally, while we compare multiple GRN inference algorithms, all evaluated methods retain an edge-centric view of network inference. Future work could extend this framework by modeling regulatory modules, temporal trajectories, perturbation responses, or dynamic TF activity states. Such extensions may help clarify when expression data provides information beyond prior refinement and how sequence-derived priors can be combined with multimodal measurements in a more flexible inference framework.

In summary, this work positions GLM-Prior as a benchmarked workflow for transferable, sequence-derived prior construction in GRN inference. Across the evaluated contexts, GLM-Prior provides a scalable strategy for constructing TF-gene priors when matched experimental assays are unavailable, with strongest support for transfer across related mammalian systems. More broadly, our results show that prior quality strongly shapes benchmarked GRN performance and that expression-based inference methods primarily provide prior-dependent refinement. These findings shift emphasis in GRN modeling toward the construction, evaluation, and transferability of prior knowledge, and provide a foundation for future methods that integrate sequence-derived priors with expression and other regulatory modalities.

## Online Methods

### The GLM-Prior model

We developed GLM-Prior, a genomic language model fine-tuned to score transcription factor (TF)-target gene pairs from nucleotide sequence and construct TF-gene prior knowledge matrices for downstream inference. Specifically, we fine-tuned the 250 million parameter Nucleotide-Transformer model (17), originally pretrained on the genomes of 850 species, to predict TF-target gene interaction probabilities from nucleotide sequences. The resulting interactions are used to define a sequence-derived prior knowledge matrix, which provides structural information for downstream GRN inference.

The model uses a transformer encoder architecture to process two distinct sequence types, TF motif sequences from the CisBP database (22) and gene body sequences derived from genome annotations in a GTF file. We append a classification head to the transformer encoder to estimate the probability that each TF-gene pair belongs to the annotaed positive class. Positive labels are derived from curated regulatory interaction resources such as STRING (23; 24; 25; 26) and TRRUST (27; 28), whereas TF-gene pairs without an annotated interaction are treated as unlabeled negative examples for supervised training. These unlabeled pairs should not be interpreted as experimentally verified non-interactions, since some may correspond to regulatory interactions that are absent from current databases or active only in specific cellular context. This design allows the model to use sequence information together with database-derived labels to construct prior knowledge matrices whose performance can be benchmarked against held-out reference networks.

Each training example is constructed by concatenating the nucleotide sequence of a TF with that of its candidate target gene, separated by a special classification token (<cls>). This composite sequence is tokenized, during which a model-specific (<cls>) token is automatically prepended to the input. The tokenized sequence is passed through the pre-trained transformer, which produces contextual embeddings across the full input. The embedding corresponding to the prepended (<cls>) token is fed into the classification head composed of a dropout layer, a linear projection, a tanh activation, a second dropout, and a final linear layer that maps to two logits representing the binary interaction label. This setup allows the model to learn sequence-associated features that improve prediction of database-derived TF-gene labels.

### Dataset construction and preprocessing

To train the model, we constructed a dataset of TF-gene pairs in which annotated positive examples were derived from curated regulatory interaction databases (YEASTRACT (11), STRING (23; 24; 25; 26) and TRRUST (27; 28)), and unlabeled negative examples were sampled from the remaining TF-gene sequence pairs without an annotated interaction. These unlabeleed pairs were used as negative examples for supervised learning, but they do not represent experimentally verified non-interactions. This dataset was split into a 99% training and 1% validation set.

Given the substantial imbalance between annotated positive and unlabeled negative samples, we implemented a downsampling strategy to reduce the number of negative samples in the training set. This preserved diversity among unlabeled negative samples while preventing the model from overfitting to the negative class. Specifically, the number of retained unlabeled negative samples after downsampling can be defined as,

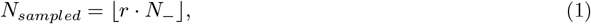

where *r* ∈ (0, 1) is the downsampling rate, and *N*_−_ is the total number of unlabeled negative examples in the dataset.

To further address class imbalance during training, we designed a custom DataLoader that performs even class batch sampling. Each batch was constructed to contain an equal number of annotated positive and unlabeled negative samples, ensuring a balanced signal during training. Specifically, each batch is defined as,

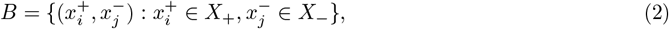

where annotated positive examples 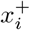 were sampled uniformly with replacement from *X*_+_ and unlabeled negative examples 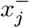 were cycled through without replacement:

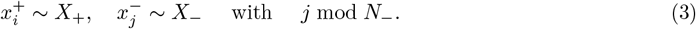

This strategy ensures that each unlabeled negative example was seen exactly once during training while annotated positive examples were reused as needed to maintain class balance. Even-class batching was critical for stabilizing training dynamics and improving the model’s sensitivity to annotated positive interactions while limiting bias toward the much larger unlabeled negative class.

### Model Architecture and Classification

The model architecture builds on the pre-trained Nucleotide Transformer by retaining its embedding and transformer encoding layers. A classification head is applied to the final hidden state corresponding to the prepended (`<cls>`) token to compute logits for a binary classification task. This classification head outputs a two-dimensional logit vector *z* ∈ ℝ^2^ representing unnormalized scores for the positive and negative classes,

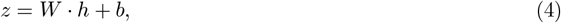

where *h* is the transformed hidden state of the (`<cls>`) token following the intermediate projection and non-linearity, and *W*, *b* are the weights and bias of the final linear layer. The components of *z* are denoted *z*_+_ and *z*_−_, corresponding to the logits for the positive and negative classes, respectively. Class probabilities are computed using a softmax function,

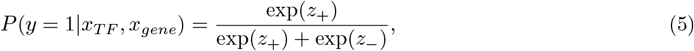

Training used a class-weighted cross-entropy loss function to account for residual class imbalance after downsampling. The loss assigns a fixed weight of 1.0 to positive examples, while the negative class weight *w*_−_ is tuned through hyperparameter search to optimize the balance between precision and recall. The loss is computed as,

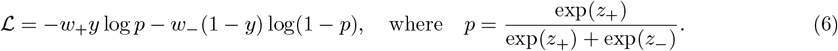

Here, *y* ∈ {0, 1} is the true label, *z*_+_ and *z*_−_ are the predicted logits for the positive and negative classes, respectively. *w*_−_ is the down-weighted negative class weight, varied during hyperparameter search, and the positive class weight *w*_+_ is fixed at 1.0. This approach maintains sensitivity to positive predictions while mitigating bias toward the negative class.

### Training procedure

We trained the model on 4 H100 GPUs using PyTorch’s Distributed Data Parallel (DDP) framework (35) to enable efficient multi-GPU scaling. The training process was distributed across GPUs to accelerate computation and ensure consistent gradient updates. We used a per-device batch size of 32 and set the gradient accumulation steps to 32, resulting in an effective batch size of 4096. The model was optimized using Adam with a learning rate of 10^−5^. Training spanned 10 epochs, using optimal hyperparameters selected through a sweep over the negative class weight (*w*_−_) and downsampling rate (see Appendix 1 for more details). We selected the configuration that achieved the highest F1 score on the validation set for final training.

After training, the model was used to score TF-gene sequence pairs without labels and construct a prior knowledge matrix for downstream GRN inference. This matrix represents sequence-derived prior evidence for putative TF-gene edges and can be used further refined using cell-type, cell-line, or condition-specific expression data during downstream GRN inference.

### Performance and Evaluation

We evaluated model performance using standard binary classification metrics, with a focus on metrics that remain robust under class imbalance. Specifically, we report precision, recall, F1 score, area under the receiver operating characteristic curve (AUC-ROC), area under the precision-recall curve (AUPRC), and Matthews correlation coefficient (MCC).

Precision and recall were computed separately for the positive and negative classes to assess the model’s ability to minimize false positives and false negatives, respectively. Let *TP*, *FP*, *FN* and *TN* denote the true positives, false positives, false negatives and true negatives. Then,

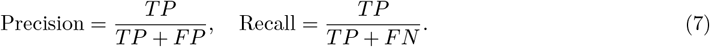

The F1 score, which represents the harmonic mean of precision and recall, was used as the primary metric for model selection and hyperparameter optimization. It can be computed as,

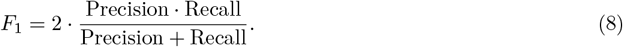

To account for class imbalance and provide a threshold-independent measure of performance, we also computed AUC-ROC and AUPRC. the ROC curve plots true positive rate (TPR) against false positive rate (FPR), defined as:

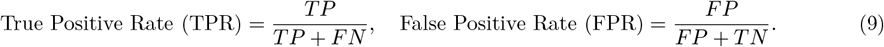

While AUC-ROC captures the model’s general discrimintative ability, AUPRC is more informative in imbalanced settings, as it directly reflects the trade-off between precision and recall. We used AUPRC to benchmark model predictions against reference networks of TF-gene interactions.

To determine the optimal classification threshold, we performed a grid search over the predicted positive class probabilities. The threshold *t*^*∗*^ that maximized the F1 score on the validation set was selected for final inference,

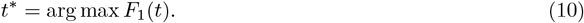

Finally, we report the Matthews correlation coefficient (MCC), a balanced measure of classification quality that incorporates all four confusion matrix components,

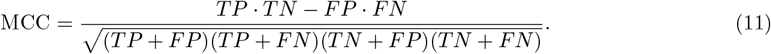

MCC ranges from −1 to +1, where +1 indicates perfect predictions, 0 indicates random predictions, and −1 indicates incorrect predictions. MCC remains informative even when classes are highly imbalanced, making it a useful complement to F1 and AUPRC.

### GRN inference with PMF-GRN

We performed GRN inference using our previously published method, PMF-GRN (Probabilistic Matrix Factorization for Gene Regulatory Network inference) (7) to estimate putative regulatory relationships between TFs and target genes from single-cell expression data. PMF-GRN is to decompose an observed gene expression matrix into latent factors that represent TF activity and regulatory interactions between TFs and their target genes. These latent factors provide an inferred, prior-conditione representation of GRN structure, which is not directly measured from gene expression data alone. Further details regarding the PMF-GRN model and the inference strategy used to obtain GRNs can be found in (6).

Using PMF-GRN, we perform inference independently on each single-cell dataset to obtain dataset-specific prior-conditioned GRNs. These inferred networks are then combined post-inference using a simple averaging strategy to produce a consensus GRN,

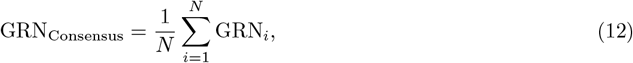

where *N* is the number of datasets and GRN_*i*_ is the inferred network for dataset *i*. This consensus GRN summarizes regulatory edges that are consistently assigned high scores across datasets, while preserving dataset-specific networks for context-specific analyses.

To evaluate the relationship between posterior uncertainty and predictive accuracy, we rank all TF-gene interaction pairs by their posterior variance, constructing 10 cumulative bins corresponding to the variances in increments of 10%. For each bin *k*, we use the posterior point estimates of the interactions within the bin and compute the AUPRC using a set of TF-gene interactions 𝒢 from a reference network. All interactions in each bin are included in the evaluation, regardless of whether they appear in the reference network.

Let ℬ_*k*_ be the set of posterior point estimates in the *k*% of variances. Then the AUPRC for bin *k* is given by:

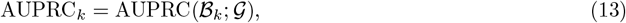

where the AUPRC is computed by comparing the predicted scores in ℬ_*k*_ to labels derived from 𝒢. The cumulative binning strategy ensures that each bin contains a sufficient number of interactions for stable AUPRC estimation.

## Declarations

## Acknowledgments

We thank the members of the Bonneau Lab and Cho Lab for insightful discussions and feedback on this manuscript. We also thank the staff of the NYU IT High Performance Computing and Flatiron Institute Scientific Computing Core. We are additionally grateful to Yanis Bahroun, Daniel Berenberg, Sarah Robinson, and Sabrina Mielke for insightful discussions related to this work.

## Author Contributions

CSG, AC, and KC jointly contributed to the conceptualization, design, and implementation of the project. CSG carried out the GLM-Prior experiments, dataset curation, validation, formal analysis, and visualizations. CSG and KC prepared the original draft. KC and RB provided supervision, project administration, and funding acquisition.

## Availability of Data and Materials

Code for GLM-Prior is available at https://github.com/cskokgibbs/GLM-Prior. Instructions on how to process nucleotide sequences for genes and TFs are available in the GLM-Prior GitHub repository, in the folder ‘`create_sequence_datasets`’. Code for previously published PMF-GRN is available at https://github.com/nyu-dl/pmf-grn.

Expression datasets used in this manuscript have the following accessions: Yeast: GSE125162 (Y1) (31) and GSE144820 (Y2) (32); Human embryonic stem cells: GSE75748 (36); Human HepG2 cells: GSE81252 (37; 38); Mouse embryonic stem cells: GSE98664 (39); Mouse dendritic Cells: GSE48968 (40); and Mouse hematopoietic stem cells: GSE81682 (41). The YEASTRACT prior knowledge was derived from the YEAS-TRACT database (11).

The prior knowledge labels used for training GLM-Prior in yeast were obtained from the YEASTRACT database (11). Prior knowledge labels for human and mouse were obtained from the STRING (23; 24; 25; 26) and TRRUST (27; 28) databases, which were provided by the BEELINE benchmark (30) and can be found at https://zenodo.org/records/7682713. Prior knowledge for Inferelator-Prior and CellOracle’s baseGRN were generated using ATAC-seq peaks from the following accessions: human embryonic stem cells: 4DNFIPGM38K4, human HepG2 cells: ENCFF913MQB, mouse embryonic stem cells: 4DNFIAEQI3RP, mouse dendritic cells: ENCFF109TUH, and mouse hematopoietic stem cells: ENCFF931CIR.

The reference network for the yeast dataset was obtained from (29). Reference networks in human embryonic stem cells, human HepG2 cells, mouse embryonic stem cells, mouse dendritic cells, and mouse hematopoietic stem cells were obtained from the BEELINE benchmarks (30).

Models for experiments from yeast, human and mouse can be found at https://huggingface.co/collections/cskokgibbs/glm-prior as well as their corresponding tokenized datasets. Priors and GRNs created with CellOracle and Inferelator software can also be found here.

## Appendix

### A. Single-Species Experiments

#### A.1 Yeast

To train our GLM-Prior model in yeast, we first obtained all 5, 999 gene body nucleotide sequences from the ENSEMBL *S. cerevisiae* (R64-1-1.UTR.gtf) genome. We obtained 212 TF sequences from the CisBP database (22) under *S. cerevisiae*.

Next, to pair these gene and TF nucleotide sequences with interaction labels, we used the YEASTRACT database of interactions (11) (6, 885 genes by 220 TFs). From these 220 YEASTRACT TFs, 46 did not have sequences associated with them from CisBP. Due to this large portion of data loss (20%), we used the promoter regions of the target genes for each missing TF as a proxy for it’s binding sequence. We defined the promoter sequence following YEASTRACT’s definition of 1000bp upstream or downstream of the gene TSS (depending on the strand orientation). Training for yeast was conducted across 5, 893 genes and 123 TFs, with 660 positive labels and 724, 179 negative labels for these interactions derived from YEASTRACT.

A hyperparameter sweep over 1 epoch of training using different class-weights and downsampling rates for the negative class revealed [0.5, 1.0] to be the optimal class-weights and 0.4 to be the optimal negative class downsampling rate. These hyperparameters were used during final training over 10 epochs. To assess robustness to model initialization, final training was repeated across three random seeds (42, 43, and 44). Validation metrics were computed on the held-out 1% of training data for each seed and are reported in Table 1. Across seeds, validation performance was stable, with ROC AUC values ranging from 0.99 to 1.00 and best F1 scores ranging from 0.76 to 0.80.

**Table 1:**
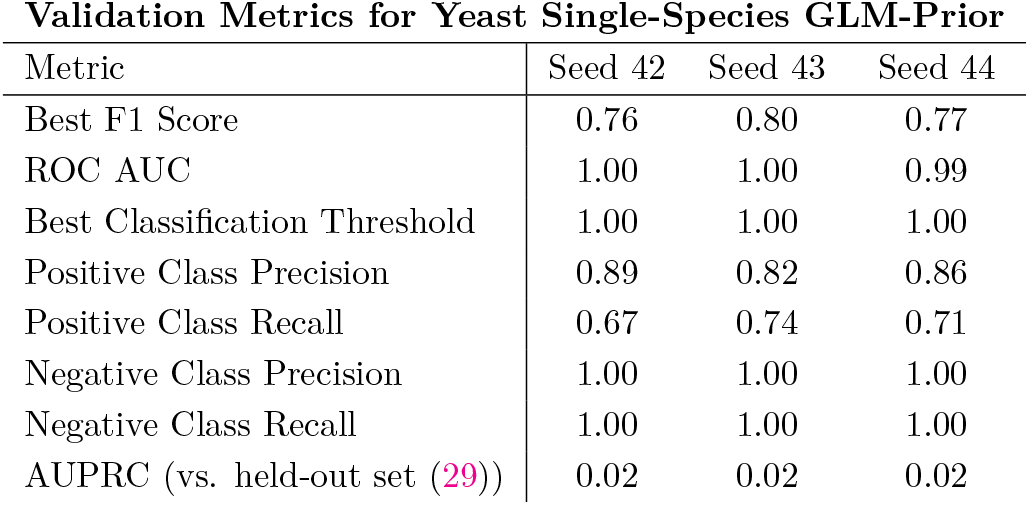
Validation performance of the GLM-Prior model trained on yeast across three random seeds. Evaluation was conducted on a held-out validation set that assesses positive and negative class contributions during training (top) and AUPRC against an independent reference network for the final inferred prior knowledge matrix (bottom).

| Validation Metrics for Yeast Single-Species GLM-Prior |  |  |  |
| --- | --- | --- | --- |
| Metric | Seed 42 | Seed 43 | Seed 44 |
| Best F1 Score | 0.76 | 0.80 | 0.77 |
| ROC AUC | 1.00 | 1.00 | 0.99 |
| Best Classification Threshold | 1.00 | 1.00 | 1.00 |
| Positive Class Precision | 0.89 | 0.82 | 0.86 |
| Positive Class Recall | 0.67 | 0.74 | 0.71 |
| Negative Class Precision | 1.00 | 1.00 | 1.00 |
| Negative Class Recall | 1.00 | 1.00 | 1.00 |
| AUPRC (vs. held-out set (29)) | 0.02 | 0.02 | 0.02 |

Following training of the yeast single-species GLM-Prior model on YEASTRACT database labels, we ran inference on all genes and TFs, including those in a held-out set not seen during training. These held-out genes and TFs corresponded to those present in the literature curated reference network from (29). Our held-out set comprised 933 genes and 98 TFs, with 986 positive labels and 95, 911 negative labels. After inference, model predictions were evaluated using AUPRC with the corresponding labels from the reference network, with a chance baseline calculated from the positive rate of the held-out set. AUPRC against this independent held-out reference was 0.02 for all three random seeds, indicating that the reported yeast prior-network performance was stable across repeated training runs.

##### A.1.1. Sequence homology experiments in yeast

Sequence homology refers to shared sequence similarity between loci, often introduced by duplication and other genome rearrangements. Because nucleotide language models are optimized to exploit recurring sequence patterns, high homology across train-test splits can, in principle, make held-out prediction easier by allowing the model to match familiar fragments rather than rely on fully generalizable features. For GLM-Prior, which is trained on labeled TF-sequence pairs, this motivates a simple check: whether genes or TFs are unusually similar to sequences seen during training, and whether controlling for that similarity changes performance at inference time. We therefore use hashFrag (42), which leverages BLAST (43) to quantify cross-split sequence similarity and generate homology-aware partitions (or prune highly similar training sequences), and then re-evaluate GLM-Prior on these homology-controlled splits to test whether sequence similarity contributes to our yeast performance.

In Figure A1A we depict the homology-aware partitioning scheme. We then quantify leakage among genes by computing, for each test gene, its maximum BLAST score to any training gene (Figure A1B-C). Using an operational cutoff *t* = 150, and holding the test set fixed (*n* = 978), pruning 346 homologous training genes reduces the fraction of “leaking” test genes from 24.8% to 7.8%.

**Figure A1:**
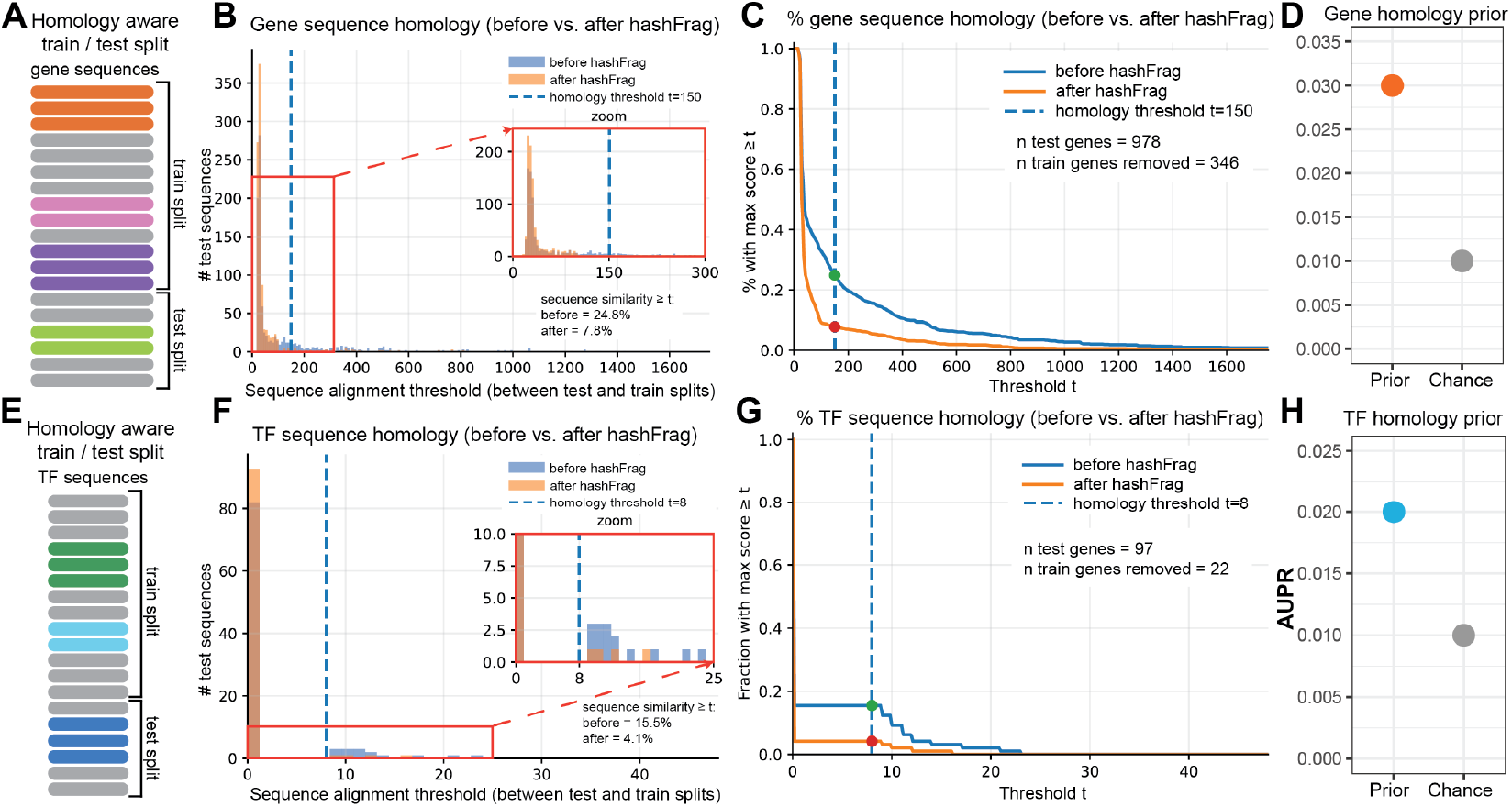
Sequence-homology analysis of yeast gene and TF splits using hashFrag. **(A)** Schematic of homology-aware partitioning, illustrating how highly similar sequences are identified and separated to reduce cross-split homology between training and test sets. **(B)** Histogram of per-test-gene maximum BLAST alignment score to any training gene, shown before (blue) and after (orange) hashFrag pruning of the training gene set. Blue dashed line indicates the similarity threshold used to define leakage. **(C)** Leakage curve for genes showing, for each similarity threshold *t*, the fraction of test genes whose maximum BLAST score to any training gene is ≥ *t*. Blue dashed line marks the leakage threshold. **(D)** AUPRC evaluation of GLM-Prior trained and tested on the hashFrag-defined gene partitions, with the gray dot indicating chance performance. **(E)** Schematic of homology-aware partitioning for TF sequences, analogous to panel A. **(F)** Histogram of per-test-TF maximum BLAST alignment score to any training TF, shown before (blue) and after (orange) hashFrag pruning of the training TF set. Blue dashed line indicates the similarity threshold. **(G)** Leakage curve for TFs showing, for each similarity threshold *t*, the fraction of test TFs whose maximum BLAST score to any training TF is ≥ *t*. Blue dashed line marks the leakage threshold. **(H)** AUPRC evaluation of GLM-Prior trained and tested on the hashFrag-defined TF partitions, with the gray dot indicating chance performance.

We next ask whether removing this homology affects model accuracy. Training on the hashFrag-defined gene partitions yields a test AUPRC of 0.03, compared to 0.01 chance performance (Figure A1D). This magnitude is comparable to the yeast results (AUPRC = 0.02) reported in Section 1, indicating that sequence leakage is not responsible for our observed performance in yeast.

We repeat the same analysis for TF sequences (Figure A1E-H). As with genes, hashFrag pruning decreases cross-split similarity, and model performance on the homology-clean TF splits remains similar to the main yeast model, again motivating against leakage-driven gains.

Overall, these experiments show that while homology-aware partitioning is methodologically important and successfully reduces cross-split similarity, it does not materially affect our model performance, which is primarily dependent on access to sufficient positive training examples. Because aggressive pruning removes substantial numbers of sequences, and thus labels, from both training and evaluation, it can reduce statistical power without measurable benefit here. For this reason, we include this analysis in the appendix to demonstrate its value, and motivate why we avoid homology pruning in the primary experiments to preserve sufficient data for training and evaluation.

#### A.2 Human cell lines (hESC & HepG2)

To train our GLM-Prior model in human, we first obtained hg38 gene body nucleotide sequences from ENSEMBL (RCh38.113.gtf). Due to the lengthy nature of human genes, and the inherent limitations of context length in large language models, we filtered our gene sequences to retain all sequences for a gene body ≤ 12, 000 nucleotides in length. This provided us with a list of 23, 533 genes. We obtained TF binding motif sequences from the CisBP database (22), under *H. sapiens*.

Training for hESC was conducted across 1, 801 genes and 504 TFs, with 5, 305 positive labels and 902, 399 negative labels for these interactions derived from STRING and TRRUST. Training for HepG2 was conducted across 2, 121 genes and 527 TFs, with 8, 087 positive labels and 1, 109, 680 negative labels for these interactions derived from STRING and TRRUST.

A hyperparameter sweep over 1 epoch of training using different class-weights and downsampling rates for the negative class revealed [0.8, 1.0] to be the optimal class-weights and 0.3 to be the optimal negative class downsampling rate in hESC, and [1.0, 1.0] to be the optimal class-weights and 0.4 to be the optimal negative class downsampling rate in HepG2. These hyperparameters were used during final training over 10 epochs. To assess robustness to model initialization, final training was repeated across three random seeds (42, 43, and 44) for each human cell line. Validation metrics were computed on held-out validation data for each seed and are reported in Table 2. Across seeds, validation performance was generally stable, although positive-class recall and best F1 score varied modestly across runs, consistent with the class imbalance of the training labels.

**Table 2:** Validation performance of the GLM-Prior model trained on human across three random seeds, evaluated in hESC and HepG2. Evaluation is conducted on held-out sets using validation metrics that consider the contribution of the positive and negative classes and AUPRC against ChIP-seq-derived reference networks after inference of the prior knowledge matrix.

| Metric | hESC |  |  | HepG2 |  |  |
| --- | --- | --- | --- | --- | --- | --- |
|  | Seed 42 | Seed 43 | Seed 44 | Seed 42 | Seed 43 | Seed 44 |
| Best F1 Score | 0.29 | 0.34 | 0.35 | 0.35 | 0.29 | 0.27 |
| ROC AUC | 0.85 | 0.81 | 0.84 | 0.83 | 0.78 | 0.80 |
| Best Classification Threshold | 0.94 | 0.99 | 0.97 | 0.97 | 0.96 | 0.92 |
| Positive Class Precision | 0.39 | 0.42 | 0.52 | 0.53 | 0.57 | 0.55 |
| Positive Class Recall | 0.22 | 0.29 | 0.26 | 0.27 | 0.18 | 0.18 |
| Negative Class Precision | 0.99 | 0.99 | 0.99 | 0.99 | 0.99 | 0.99 |
| Negative Class Recall | 1.00 | 1.00 | 1.00 | 1.00 | 1.00 | 1.00 |
| AUPRC (vs. reference network) | 0.19 | 0.20 | 0.19 | 0.30 | 0.29 | 0.30 |

Following training of the human single-species GLM-Prior model on STRING and TRRUST database labels, we ran inference on all genes and TFs, including those in a held-out set not seen during training. These held-out genes and TFs corresponded to those present in the BEELINE hESC and HepG2 reference ChIP-seq networks, respectively. For hESC, our held-out set comprised 4, 773 genes and 79 TFs, with 58, 337 positive labels, and 318, 730 negative labels. For HepG2, our held-out set comprised 4, 338 genes and 53 TFs, with 56, 567 positive labels, and 173, 347 negative labels. After inference, model predictions were evaluated using AUPRC with the corresponding labels from the hESC and HepG2 reference ChIP-seq networks, with a chance baseline calculated from the positive rate of each held-out set. AUPRC against the independent reference networks was stable across seeds, ranging from 0.19 to 0.20 in hESC and from 0.29 to 0.30 in HepG2.

#### A.3 Mouse cell lines (mESC, mDC, & mHSC)

To train our GLM-Prior model in mouse, we first obtained the mm10 gene body nucleotide sequences from ENSEMBL (GRCm39.113.gtf). Similarly to the single-species human experiments, we again filtered the length of our mouse genes to retain all sequences for a gene body 12, 000 nucleotides in length. This provided us with a list of 37, 755 genes. We obtained TF binding motif sequences from the CisBP database (22), under *M. musculus*.

Training for mESC was conducted across 1, 097 genes and 437 TFs, with 3, 129 positive labels and 476, 260 negative labels for these interactions derived from STRING and TRRUST. Training for mDC was conducted across 2, 837 genes and 462 TFs, with 11, 551 positive labels and 1, 299, 134 negative labels for these interactions derived from STRING and TRRUST. Training for mHSC was conducted across 1, 129 genes and 419 TFs, with 2, 808 positive labels and 470, 243 negative labels.

A hyperparameter sweep over 1 epoch of training using different class-weights and downsampling rates for the negative class revealed [0.2, 1.0] to be the optimal class-weights and 0.5 to be the optimal negative class downsampling rate in mESC, and [0.1, 1.0] to be the optimal class-weights and 0.4 to be the optimal negative class downsampling rate in mDC, and [0.1, 1.0] to be the optimal class-weights and 0.5 to be the optimal negative class downsampling rate in mHSC. These hyperparameters were used during final training over 10 epochs. To assess robustness to model initialization, final training was repeated across three random seeds (42, 43, and 44) for each mouse cell line. Validation metrics were computed on held-out validation data for each seed and are reported in Table 3. Across seeds, validation metrics were broadly consistent, although positive-class precision, positive-class recall, and best F1 score showed some variation, reflecting the difficulty of learning from highly imbalanced regulatory labels.

**Table 3:** Validation performance of the GLM-Prior model trained on mouse across three random seeds, evaluated in mESC, mDC, and mHSC. Evaluation is conducted on held-out sets using validation metrics that consider the contribution of the positive and negative classes and AUPRC against ChIP-seq-derived reference networks after inference of the prior knowledge matrix.

| Validation Metrics for Mouse Single-Species GLM-Prior |  |  |  |  |  |  |  |  |  |
| --- | --- | --- | --- | --- | --- | --- | --- | --- | --- |
| Metric | mESC |  |  | mDC |  |  | mHSC |  |  |
|  | Seed 42 | Seed 43 | Seed 44 | Seed 42 | Seed 43 | Seed 44 | Seed 42 | Seed 43 | Seed 44 |
| Best F1 Score | 0.19 | 0.15 | 0.16 | 0.20 | 0.24 | 0.17 | 0.19 | 0.22 | 0.30 |
| ROC AUC | 0.70 | 0.68 | 0.74 | 0.80 | 0.75 | 0.75 | 0.74 | 0.68 | 0.75 |
| Best Classification Threshold | 0.97 | 0.97 | 0.97 | 0.99 | 0.99 | 0.99 | 0.98 | 0.94 | 0.90 |
| Positive Class Precision | 0.23 | 0.24 | 0.23 | 0.17 | 0.18 | 0.35 | 0.50 | 0.33 | 0.46 |
| Positive Class Recall | 0.16 | 0.11 | 0.12 | 0.19 | 0.21 | 0.11 | 0.12 | 0.16 | 0.23 |
| Negative Class Precision | 0.99 | 0.99 | 0.99 | 0.99 | 0.99 | 0.99 | 0.99 | 0.99 | 0.99 |
| Negative Class Recall | 0.99 | 1.00 | 1.00 | 0.99 | 0.99 | 1.00 | 1.00 | 1.00 | 1.00 |
| AUPRC (vs. reference network) | 0.23 | 0.24 | 0.22 | 0.09 | 0.09 | 0.09 | 0.49 | 0.47 | 0.44 |

**Table 4:** Statistical comparison of gene body sequence properties between training and test gene sets across six cell lines. Gene length (bp) and GC fraction are compared using the Kolmogorov-Smirnov (KS) statistic and *p*-value.

| Cell line | Feature | Mean |  | Median |  | KS test |  |
| --- | --- | --- | --- | --- | --- | --- | --- |
| | | Train | Test | Train | Test | stat | $p$ |
| Yeast | Length (bp) | 1681.9 | 1445.6 | 1395.0 | 1308.0 | 0.089 | $1.41 \times 10^{-6}$ |
| | GC fraction | 0.389 | 0.394 | 0.385 | 0.392 | 0.115 | $1.20 \times 10^{-10}$ |
| hESC | Length (bp) | 2403.7 | 5452.2 | 704.0 | 5203.0 | 0.471 | $5.33 \times 10^{-264}$ |
| | GC fraction | 0.488 | 0.532 | 0.482 | 0.538 | 0.234 | $2.68 \times 10^{-63}$ |
| HepG2 | Length (bp) | 2934.1 | 5482.9 | 1275.0 | 5271.5 | 0.354 | $5.37 \times 10^{-159}$ |
| | GC fraction | 0.500 | 0.530 | 0.496 | 0.534 | 0.163 | $2.18 \times 10^{-33}$ |
| mESC | Length (bp) | 2886.8 | 5803.2 | 1058.5 | 5546.0 | 0.444 | $1.01 \times 10^{-156}$ |
| | GC fraction | 0.478 | 0.503 | 0.473 | 0.506 | 0.220 | $1.18 \times 10^{-37}$ |
| mDC | Length (bp) | 4373.9 | 6089.4 | 3775.0 | 5910.0 | 0.228 | $2.00 \times 10^{-69}$ |
| | GC fraction | 0.500 | 0.502 | 0.503 | 0.505 | 0.044 | $5.87 \times 10^{-3}$ |
| mHSC | Length (bp) | 2357.7 | 5445.2 | 136.0 | 5077.0 | 0.535 | $5.61 \times 10^{-237}$ |
| | GC fraction | 0.488 | 0.494 | 0.479 | 0.497 | 0.106 | $2.93 \times 10^{-9}$ |

**Table 5:** Performance of the gene k-mer composition baseline on held-out test pairs across six cell lines. A Ridge regression model was trained to predict each gene’s positive interaction rate from 4-mer body composition using training genes only, and the resulting gene-level scores were broadcast to all test pairs sharing that gene. AUPRC and AUROC are reported alongside the positive-rate baseline for each cell line. AUROC near 0.50 across all cell lines indicates near-chance ranking of positive interactions despite modest AUPRC values driven by differences in reference network label density.

| Gene $k$-mer composition baseline: interaction label prediction | | | |
| --- | --- | --- | --- |
| Cell line | Positive-rate baseline | AUPRC | AUROC |
| Yeast | 0.01 | 0.02 | 0.53 |
| hESC | 0.15 | 0.16 | 0.52 |
| HepG2 | 0.25 | 0.26 | 0.52 |
| mESC | 0.20 | 0.21 | 0.52 |
| mDC | 0.09 | 0.09 | 0.49 |
| mHSC | 0.37 | 0.43 | 0.55 |

**Table 6:** Performance of the annotation degree baseline on held-out test pairs across six cell lines. A logistic regression classifier was trained on gene- and TF-level annotation degree features from the training prior, capturing how frequently each gene and TF appeared as a positive interaction in STRING, TRRUST, or YEASTRACT, and applied to held-out test pairs. AUROC of 0.50 in all cell lines indicates that training annotation frequency has no predictive relationship to test reference network labels.

| Annotation degree baseline: interaction label prediction |  |
| --- | --- |
| Cell line | AUROC |
| Yeast | 0.50 |
| hESC | 0.50 |
| HepG2 | 0.50 |
| mESC | 0.50 |
| mDC | 0.50 |
| mHSC | 0.50 |

Following training of the mouse single-species GLM-Prior model on STRING and TRRUST database labels, we ran inference on all genes and TFs, including those in a held-out set not seen during training. These held-out genes and TFs corresponded to those present in the BEELINE mESC, mDC and mHSC reference ChIP-seq networks, respectively. For mESC, our held-out set comprised 5, 711 genes and 59 TFs, with 65, 933 positive labels, and 271, 016 negative labels. For mDC, our held-out set comprised 3, 373 genes and 29 TFs, with 8, 708 positive labels, and 89, 109 negative labels. For mHSC, our held-out set comprised 6, 777 genes and 72 TFs, with 182, 005 positive labels, and 305, 939 negative labels. After inference, model predictions were evaluated using AUPRC with the corresponding labels from the mESC, mDC, and mHSC reference ChIP-seq networks, with a chance baseline calculated from the positive rate of each held-out set. AUPRC against the independent reference networks was stable in mESC (0.22-0.24) and mDC (0.09 across all seeds), and showed modest variation in mHSC (0.44-0.49).

#### A.4 Diagnostic analyses of gene body composition and database annotation features

To assess whether GLM-Prior predictions could be explained by non-regulatory proxy features, we performed a series of diagnostic analyses across all six cell lines. These analyses tested whether model performance could arise from gene body sequence composition, gene length, GC content, or database annotation patterns, rather than from regulatory sequence information. Where proxy models were evaluated for regulatory interaction prediction, they were trained exclusively on training data and evaluated on held-out test pairs without access to test interaction labels, matching the evaluation setting used for GLM-Prior.

We first asked whether training and test set genes differ systematically in coarse gene body sequence properties. Gene body length and GC content were quantified for each gene and compared between training and test splits using nonparametric distributional tests. Test genes tended to be longer and more GC-rich than training genes, with statistically significant differences across the mammalian cell lines and more modest distributional differences in yeast (Supplementary Figure A2A-D and Supplementary Table 4). To determine whether these properties were sufficient to distinguish the two splits, we trained a logistic regression classifier to predict train versus test membership using only gene length and GC content. This classifier achieved AUROC values ranging from 0.57 in yeast to 0.84 in mESC (Supplementary Figure A3), indicating that the splits are partially separable by coarse gene body features.

We then repeated this analysis using gene body *k*-mer composition. Classifiers trained on 4-, 5-, and 6-mer profiles also distinguished training from test genes, with 4-mer AUROC ranging from 0.68 in yeast to 0.84 in mESC and declining at longer *k*-mer sizes in five of six cell lines (Supplementary Figure A4A). The gene-disjoint split therefore retains measurable compositional differences between training and test genes, providing a stringent setting in which to test whether such features could account for GLM-Prior’s performance. If gene length, GC content, or local *k*-mer composition were responsible, models trained directly on these features would recover held-out regulatory labels, and would do so under the same train-test distributional shift faced by GLM-Prior.

**Figure A2:**
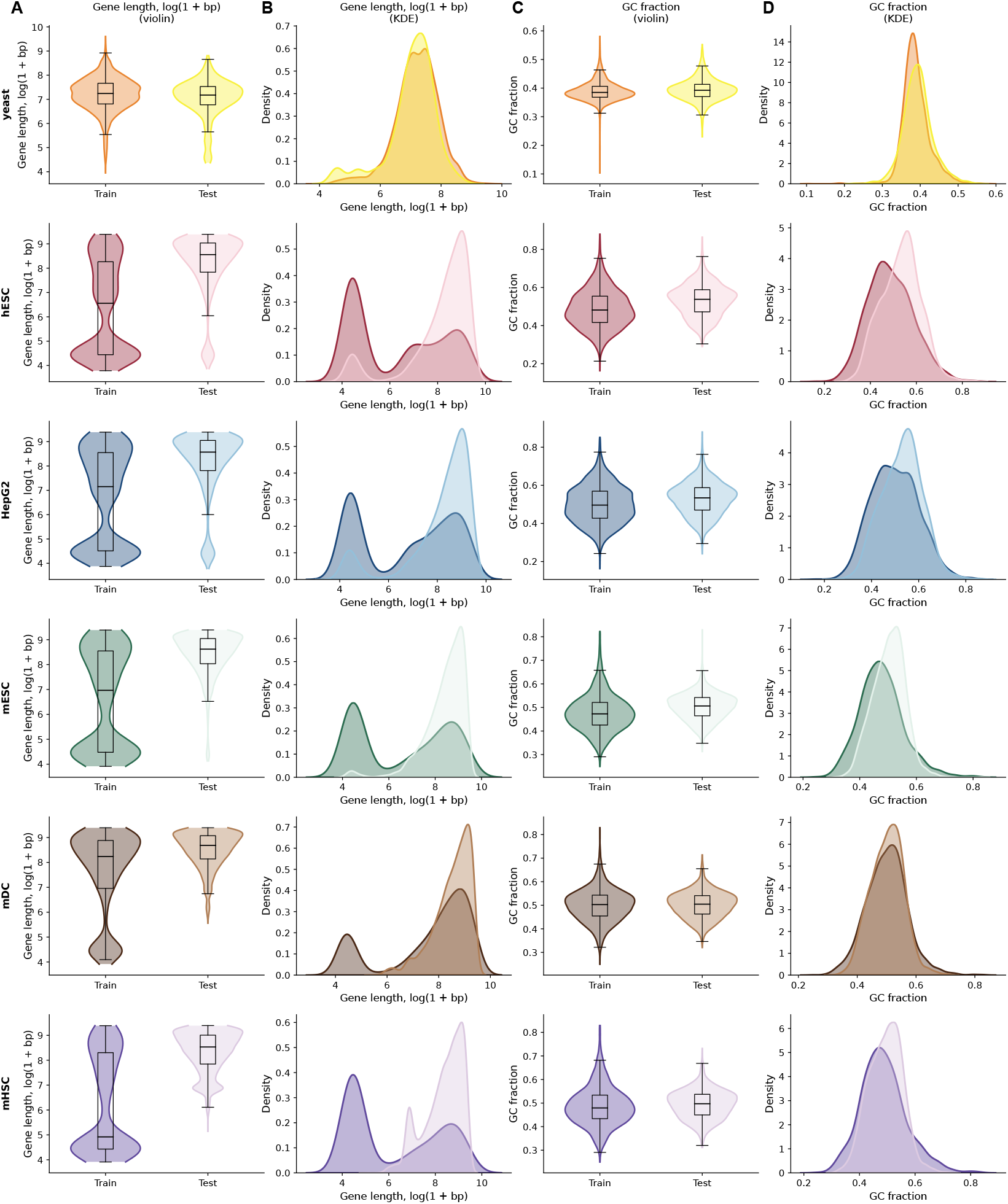
Comparison of gene body sequence properties between training and test gene sets across six cell lines. Violin plots and kernel density estimates show the distribution of gene body length (log(1 + bp), panels **A** and **B**) and GC fraction (panels **C** and **D**) for training and test genes in each cell line. Test genes are consistently longer and more GC-rich than training genes across mammalian cell lines, while yeast shows comparatively modest distributional differences between splits.

**Figure A3:**
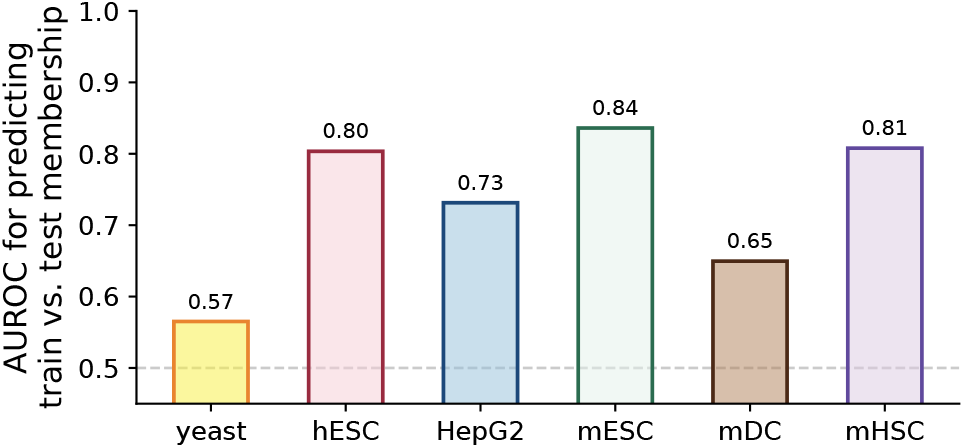
AUROC for a logistic regression classifier trained to distinguish training from test genes using basic sequence features (gene body length and GC content) across six cell lines. AUROC above 0.50 indicates that training and test genes are separable by these coarse sequence properties alone, with values ranging from 0.57 in yeast to 0.84 in mESC. The dashed line indicates chance performance (AUROC = 0.50).

**Figure A4:**
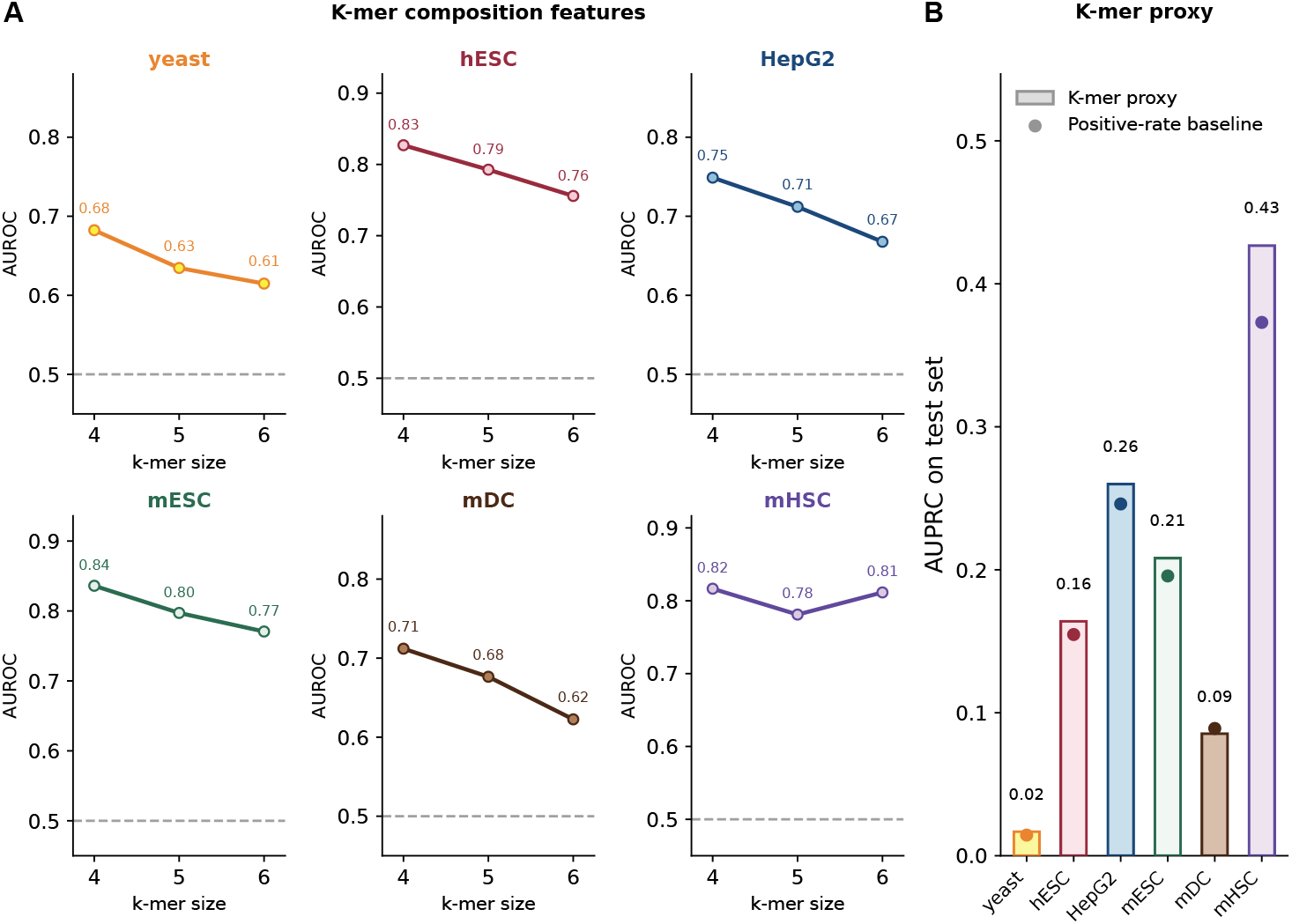
Gene body *k*-mer composition distinguishes training from test genes but has limited predictive value for held-out regulatory interaction labels. (**A**) AUROC for logistic regression classifiers trained to distinguish training from test genes using gene body *k*-mer composition features (*k* = 4, 5, and 6) across six cell lines. AUROC above 0.50 indicates that training and test genes are compositionally separable at the corresponding *k*-mer resolution. (**B**) AUPRC of a gene *k*-mer composition proxy baseline evaluated on held-out test pairs. A Ridge regression model was trained using training genes only to predict each gene’s positive interaction rate from 4-mer body composition, and the resulting gene-level scores were broadcast to all test pairs sharing that gene. Bars show AUPRC and dots indicate the positive-rate baseline for each cell line.

We first assessed whether gene body *k*-mer composition could predict held-out regulatory interaction labels. A gene *k*-mer composition baseline was trained to predict each gene’s regulatory propensity from 4-mer body composition using training genes only, and the resulting gene-level scores were evaluated on held-out test pairs. This baseline achieved AUPRC near or below the positive-rate baseline in most cell lines (Supplementary Figure A4B) and AUROC values near 0.50 across all cell lines (range: 0.49 to 0.55; Supplementary Table 5), indicating near-chance ranking of positive and negative regulatory interactions.

We next evaluated whether broader sequence-derived proxy baselines could explain held-out regulatory interaction performance. We considered three baselines: basic gene-body features, gene *k*-mer composition, and a combined sequence baseline incorporating both coarse and compositional features (Supplementary Figure A5). A classifier trained on basic gene body features, including sequence length and GC content, achieved AUPRC at or below the positive-rate baseline in five of six cell lines (yeast: 0.02 vs. 0.01 baseline; hESC: 0.12 vs. 0.15; mESC: 0.17 vs. 0.20; mDC: 0.08 vs. 0.09; mHSC: 0.36 vs. 0.37), with only a modest gain in HepG2 (0.27 vs. 0.25). The combined sequence baseline achieved AUPRC at or below the positive-rate baseline in yeast (0.01 vs. 0.01) and mDC (0.09 vs. 0.09), with modest above-baseline performance in hESC (0.17 vs. 0.15), HepG2 (0.28 vs. 0.25), mESC (0.22 vs. 0.20), and mHSC (0.42 vs. 0.37). In the same comparison, GLM-Prior matched or exceeded this combined sequence baseline across all six cell lines (yeast: 0.02 vs. 0.01; hESC: 0.19 vs. 0.17; HepG2: 0.30 vs. 0.28; mESC: 0.23 vs. 0.22; mDC: 0.09 vs. 0.09; mHSC: 0.49 vs. 0.42). Thus, although gene body properties carry limited signal in some mammalian reference networks, they do not account for GLM-Prior’s predictive performance.

Finally, we assessed whether database annotation patterns could explain test performance. We trained a classifier using gene- and TF-level annotation degree features from the training prior, capturing how frequently each gene and TF appeared as a positive interaction in STRING, TRRUST, or YEASTRACT. This baseline tests whether genes or TFs that are more frequently annotated in the training database are also more likely to appear as positives in the held-out reference networks. Although these degree features showed strong predictive ability within the training annotation matrix, they achieved AUROC of exactly 0.50 in all six held-out test cell lines (Supplementary Table 6). Training annotation frequency therefore has no detectable predictive relationship with held-out test reference labels.

Together, these analyses show that the train and test gene sets differ detectably in gene body length, GC content, and *k*-mer composition, establishing a plausible axis along which proxy learning could occur. However, direct proxy baselines trained on these same features show limited and inconsistent ability to predict held-out regulatory interaction labels, with near-chance AUROC for *k*-mer-derived gene propensities and no predictive value from annotation-degree features. The compositional separation between train and test genes therefore does not explain GLM-Prior’s held-out performance. GLM-Prior instead generalizes across a nontrivial gene-level distributional shift while performing at or above baselines that explicitly encode the proxy features tested here. These results indicate that GLM-Prior’s predictions are not reducible to coarse gene body composition or database annotation frequency, while remaining consistent with the conclusion that the broader finding that the model improves agreement with available regulatory reference annotations.

**Figure A5:**
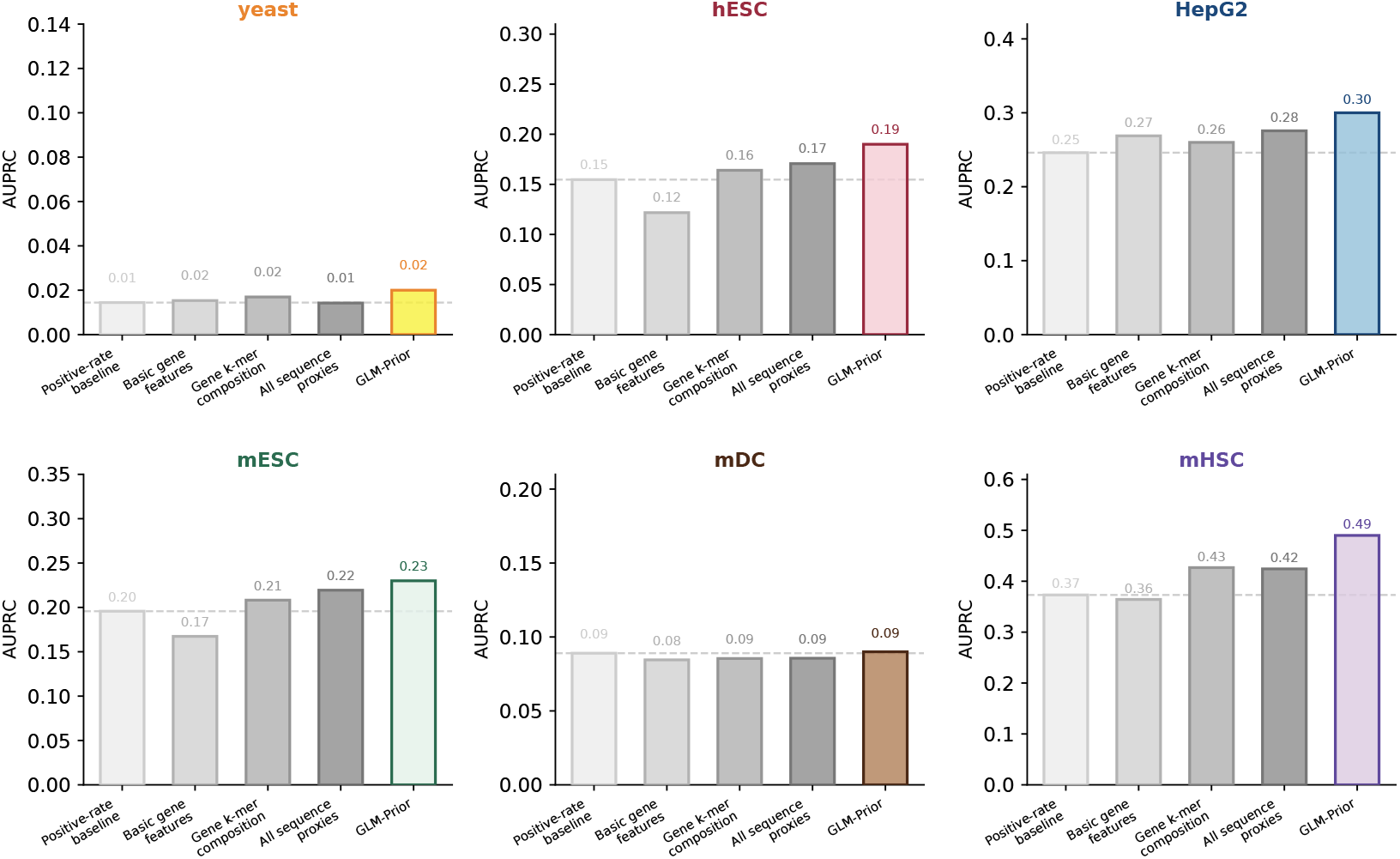
Comparison of sequence-derived baselines and GLM-Prior AUPRC on held-out test pairs across six cell lines. Each panel shows four gray bars: the positive-rate baseline, a basic gene-body features baseline (sequence length and GC content), a gene k-mer composition baseline (4-mer body composition), and a combined baseline incorporating both sequence length, GC content, and k-mer-derived propensity scores. GLM-Prior AUPRC is shown in the cell line color for each panel. GLM-Prior matches or exceeds the combined sequence baseline in all six cell lines, demonstrating that its predictive performance is not accounted for by gene body sequence composition alone. The dashed line indicates the positive-rate baseline for each cell line.

#### A.5 External support for high-confidence GLM-Prior predictions in HepG2

To provide an additional biological evaluation of high-confidence GLM-Prior predictions, we examined candidate TF-gene interactions in HepG2 that were absent from the held-out BEELINE reference network but supported by independent evidence sources. Specifically, we focused on BEELINE-negative test-set edges that were ranked within the top 1% of GLM-Prior predictions and were also supported by STRING and TRRUST annotations. These STRING and TRRUST annotations correspond to interactions that were excluded from model training because they overlapped with genes or TFs in the BEELINE test set. Therefore, although they provide independent database support, these specific labels were not observed by the model during training. This analysis asks whether high-confidence GLM-Prior predictions include plausible regulatory interactions beyond those captured in the BEELINE reference network, providing an additional literature- and database-supported assessment of candidate edges.

**Figure A6:**
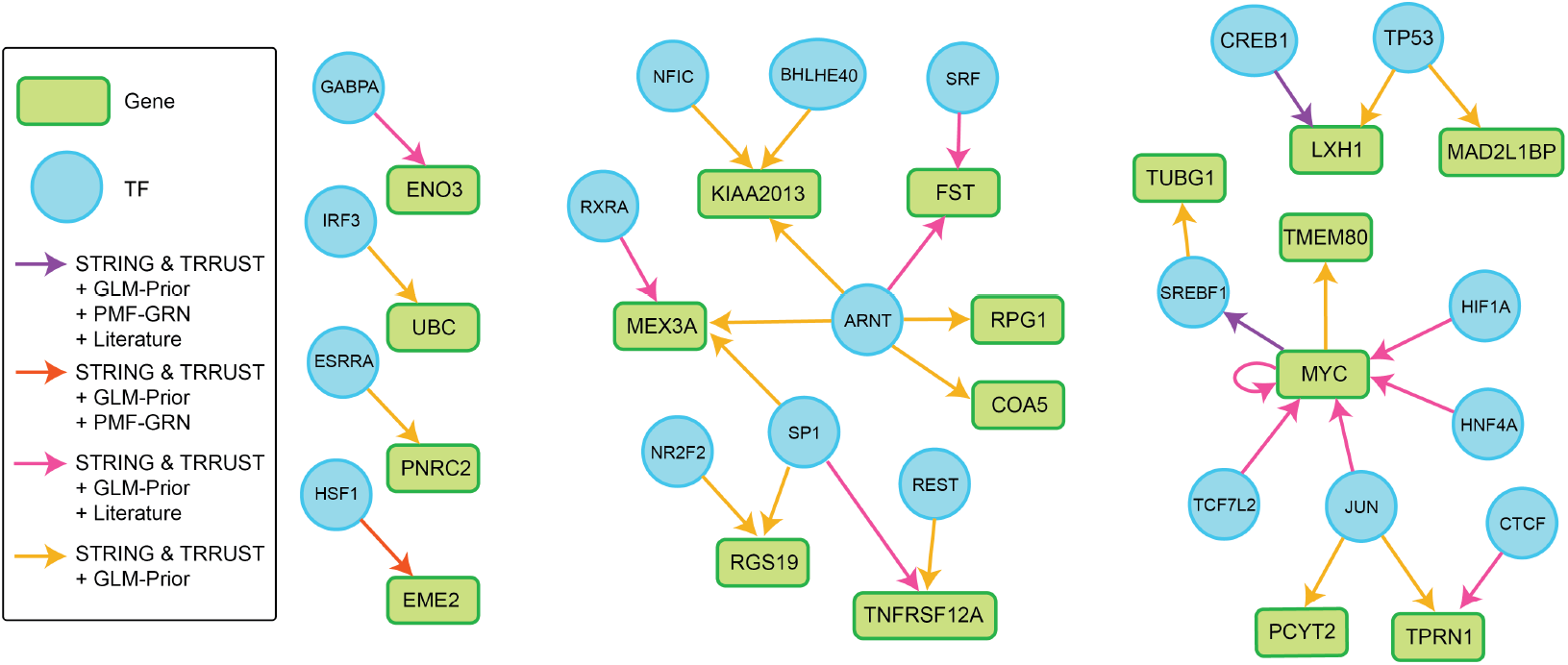
Network visualization of candidate HepG2 TF-gene interactions identified among BEELINE-negative held-out test edges. Green boxes indicate genes and blue circles indicate transcription factors. Directed edges indicate candidate regulatory interactions supported by GLM-Prior and held-out STRING and TRRUST annotations, with edge color denoting the combination of supporting evidence. Purple arrows indicate interactions supported by STRING and TRRUST, GLM-Prior, PMF-GRN, and literature review; orange arrows indicate interactions supported by STRING and TRRUST, GLM-Prior, and PMF-GRN; pink arrows indicate interactions supported by STRING and TRRUST, GLM-Prior, and literature review; and yellow arrows indicate interactions supported by STRING and TRRUST and GLM-Prior.

Using this criterion, we identified 32 BEELINE-negative edges that were supported by both GLM-Prior and held-out STRING/TRRUST annotations (Figure A6, Table 7). Among these, three interactions were also supported by PMF-GRN: HSF1 → EME2, SREBF1 → MYC, and CREB1 → LHX1. These three edges represent the strongest level of agreement in this analysis, as they are supported by external database annotations, prioritized by the sequence-derived GLM-Prior model, and retained after expression-based GRN inference with PMF-GRN. Of these three jointly supported edges, CREB1 → LHX1 and SREBF1 → MYC were also supported by literature review, providing an example in which held-out database support, GLM-Prior, PMF-GRN, and published regulatory evidence all agree.

**Table 7:** Candidate TF-gene interactions predicted by GLM-Prior among BEELINE-negative held-out test edges. All listed edges are absent from the BEELINE held-out reference network but are supported by STRING and TRRUST annotations and GLM-Prior. Additional support from PMF-GRN and literature is indicated where available.

| Gene | TF | STRING/TRRUST | GLM-Prior | PMF-GRN | Literature |
| --- | --- | --- | --- | --- | --- |
| EME2 | HSF1 | ✓ | ✓ | ✓ | — |
| MYC | SREBF1 | ✓ | ✓ | ✓ | ✓ (44; 45) |
| LHX1 | CREB1 | ✓ | ✓ | ✓ | ✓ (46) |
| TNFRSF12A | SP1 | ✓ | ✓ | — | ✓ (47) |
| UBC | IRF3 | ✓ | ✓ | — | — |
| TNFRSF12A | REST | ✓ | ✓ | — | — |
| MEX3A | SP1 | ✓ | ✓ | — | — |
| MEX3A | RXRA | ✓ | ✓ | — | ✓ (48) |
| MEX3A | ARNT | ✓ | ✓ | — | — |
| KIAA2013 | NFIC | ✓ | ✓ | — | — |
| KIAA2013 | ARNT | ✓ | ✓ | — | — |
| MYC | JUN | ✓ | ✓ | — | ✓ (49; 50) |
| KIAA2013 | BHLHE40 | ✓ | ✓ | — | — |
| MYC | HIF1A | ✓ | ✓ | — | ✓ (51; 52; 53) |
| MYC | MYC | ✓ | ✓ | — | ✓ (54; 55) |
| TMEM80 | MYC | ✓ | ✓ | — | — |
| TRNP1 | JUN | ✓ | ✓ | — | — |
| ENO3 | GABPA | ✓ | ✓ | — | ✓ (56; 48) |
| RGP1 | ARNT | ✓ | ✓ | — | — |
| MYC | HNF4A | ✓ | ✓ | — | ✓ (57) |
| MYC | TCF7L2 | ✓ | ✓ | — | ✓ (58; 59) |
| MAD2L1BP | TP53 | ✓ | ✓ | — | — |
| PNRC2 | ESRRA | ✓ | ✓ | — | — |
| COA5 | ARNT | ✓ | ✓ | — | — |
| RGS19 | SP1 | ✓ | ✓ | — | — |
| RGS19 | NR2F2 | ✓ | ✓ | — | — |
| FST | SRF | ✓ | ✓ | — | ✓ (60; 61) |
| TRNP1 | CTCF | ✓ | ✓ | — | ✓ (62) |
| FST | ARNT | ✓ | ✓ | — | ✓ (63) |
| PCYT2 | JUN | ✓ | ✓ | — | — |
| LHX1 | TP53 | ✓ | ✓ | — | — |
| TUBG1 | SREBF1 | ✓ | ✓ | — | — |

A literature review identified additional candidate interactions with published support, including examples supported by promoter binding, curated transcription factor target annotations, perturbation-associated expression changes, or functional regulatory relationships. These included REST → TNFRSF12A, RXRA → MEX3A, JUN → MYC, HIF1A → MYC, MYC autoregulation, GABPA → ENO3, HNF4A → MYC, TCF7L2 → MYC, SRF → FST, and ARNT → FST. In several cases, the supporting evidence was not HepG2-specific, which is consistent with the cell-line-agnostic nature of the GLM-Prior training labels. In other cases, the evidence supported regulation through a broader TF complex or pathway rather than direct regulation by the individual TF alone. We therefore treat these examples as biologically plausible candidate interactions rather than definitive HepG2-specific regulatory edges.

This analysis is not intended as a comprehensive experimental validation of newly inferred GLM-Prior edges. Instead, it provides a targeted qualitative assessment of whether high-confidence GLM-Prior predictions that are absent from the BEELINE held-out reference may correspond to interactions supported by other regulatory resources or published studies. The presence of held-out STRING/TRRUST support, PMF-GRN agreement for a subset of edges, and literature evidence for several BEELINE-negative predictions is consistent with the incompleteness of benchmark reference networks and suggests that some GLM-Prior predictions labeled as false positives relative to BEELINE may represent plausible missing annotations. Because many literature-supported cases are context-dependent, indirect, or derived from non-HepG2 systems, these results are best interpreted as hypothesis-generating examples that complement the benchmark-based evaluation.

### B. Transfer-Learning Experiments

Transfer learning experiments were conducted by first ensuring that no overlap existed between the training set (human, mouse, or yeast), and the corresponding inference set with held-out evaluation labels. For the human and mouse model that transferring knowledge to yeast, no information was overlapping between training and inference *a priori*.

For the mouse trained model which transferred knowledge to hESC and HepG2, two separate models were trained, one in which overlaps between the mouse genes, TFs and labels with hESC were removed, and the second in which the overlaps between mouse genes, TFs and labels with HepG2 were removed. For the human trained model which transferred knowledge to mESC, mDC, and mHSC, three separate models were trained. In the first model, overlaps between human genes, TFs, and labels with mESC were removed. In the second model, overlaps between human genes, TFs, and labels with mDC were removed. In the third model, overlaps between human genes, TFs, and labels with mHSC were removed.

Training six separate models allowed us to ensure an optimal number of gene and TF sequences and their corresponding labels were seen during training, while maintaining complete independence from the inference set and corresponding evaluation performance. Results for the transfer learning experiments can be found in Table 8.

**Table 8:** AUPRC scores for cross-species transfer learning. Each row indicates the species used to train the GLM-Prior model and the target species on which GRN inference was performed. Evaluation was conducted against species-specific ChIP-seq or curated reference datasets.

| AUPRC Results from Transfer Learning |  |  |
| --- | --- | --- |
| Training Dataset | Inference Dataset | AUPRC |
| Human and Mouse | Yeast | 0.02 |
| Mouse | hESC | 0.19 |
| Mouse | HepG2 | 0.31 |
| Human | mESC | 0.23 |
| Human | mDC | 0.09 |
| Human | mHSC | 0.46 |

### C Multi-Species Experiments

To train a multi-species model, we consecutively trained GLM-Prior on human, mouse, and yeast. To ensure no sequence overlap between the genes, TFs, and labels used during training and inference, we removed all overlapping inference sets from each individual organisms training set. Here, we ensured there was no inter-species leakage, as well as intra-species leakage, between training and inference sets.

Following the consecutive training of the multi-species model, we ran inference in yeast, hESC, HepG2, mESC, mDC, and mHSC held-out sets, respectively. Results for this multi-species model inference can be found in Table 9.

**Table 9:** AUPRC scores for multi-species model inference in human, mouse, and yeast. GRNs were inferred using a unified model trained jointly on all three species, and evaluated against cell-line specific reference networks.

| AUPRC Results from Multi-Species Training |  |  |
| --- | --- | --- |
| Training Dataset | Inference Dataset | AUPRC |
| Human + Mouse + Yeast | Yeast | 0.02 |
|  | hESC | 0.18 |
|  | HepG2 | 0.29 |
|  | mESC | 0.23 |
|  | mDC | 0.09 |
|  | mHSC | 0.49 |

### D Inferelator-Prior and CellOracle baseGRN

We generated prior knowledge for Inferelator-Prior (6) and CellOracle’s baseGRN (8), using each respective methods published Python software. To generate prior knowledge with Inferelator-Prior and CellOracle baseGRN for each of the six cell lines, we obtained ATAC-seq datasets from the following accessions: Human embryonic stem cells: 4DNFIPGM38K4, Human HepG2 cells: ENCFF913MQB, Mouse embryonic stem cells: 4DNFIAEQI3RP, Mouse dendritic cells: ENCFF109TUH, and Mouse hematopoietic stem cells: ENCFF931CIR. In yeast, prior knowledge datasets were obtained from (6) without further modification.

These priors were constructed using motifs from CisBP, the same motifs used in GLM-Prior to ensure compatibility and fairness during evaluation.

Results for the Inferelator-Prior and CellOracle baseGRN prior knowledge for each six cell lines using the same reference sets described in Section A are provided in Table 10.

**Table 10.** AUPRC of GLM-Prior, Inferelator-Prior and CellOracle baseGRN across six cell lines.

| Cell Line | AUPRC |  |  |
| --- | --- | --- | --- |
|  | GLM-Prior | Inferelator-Prior | CellOracle baseGRN |
| Yeast | 0.02 | 0.03 | 0.05 |
| hESC | 0.19 | 0.17 | 0.16 |
| HepG2 | 0.30 | 0.24 | 0.36 |
| mESC | 0.23 | 0.17 | 0.17 |
| mDC | 0.09 | 0.07 | 0.07 |
| mHSC | 0.49 | 0.34 | 0.34 |

### E GRN Inference with PMF-GRN, the Inferelator, and CellOracle

GRN inference for each of the six cell lines was performed using PMF-GRN, the Inferelator 3.0, and CellOracle Python software, respectively.

Single cell gene expression datasets were obtained from GSE125162 (38, 225 cells by 6, 763 genes) (31) and GSE144820 (6, 118 cells by 6, 763 genes) (32) without modification. The dataset was then combined by concatenating GSE125162 and GSE144820 on the cells axis (44, 343 cells by 6, 763 genes). The remaining hESC (758 cells by 17, 735 genes), HepG2 (425 cells by 11, 515 genes), mESC (421 cells by 18, 385 genes), mDC (383 cells by 7, 371 genes), and mHSC (2, 807 cells by 4, 762 genes) single-cell expression datasets were obtained from BEELINE (30).

Prior knowledge for each method is required to produce a GRN from single-cell expression data. For the experiments in this section, we use the specific cell line prior knowledge generated by the corresponding algorithm (described in Section A for GLM-Prior (using seed 42), and Section 10 for Inferelator-Prior and CellOracle baseGRN. Results for these GRN inference experiments can be found in Table 11).

**Table 11:** AUPRC of three GRN inference methods (PMF-GRN, Inferelator 3.0, and CellOracle) across six cell lines.

| Cell Line | AUPRC |  |  |
| --- | --- | --- | --- |
|  | PMF-GRN | Inferelator 3.0 | CellOracle |
| Yeast | 0.02 | 0.04 | 0.06 |
| hESC | 0.19 | 0.20 | 0.17 |
| HepG2 | 0.38 | 0.36 | 0.38 |
| mESC | 0.27 | 0.25 | 0.37 |
| mDC | 0.10 | 0.09 | 0.09 |
| mHSC | 0.52 | 0.42 | 0.39 |

Additional metrics for GLM-Prior and PMF-GRN, including F1 score, precision, recall, ROC-AUC, positive class recall, positive class precision, negative class recall, and negative class precision, are provided in A7.

**Figure A7:**
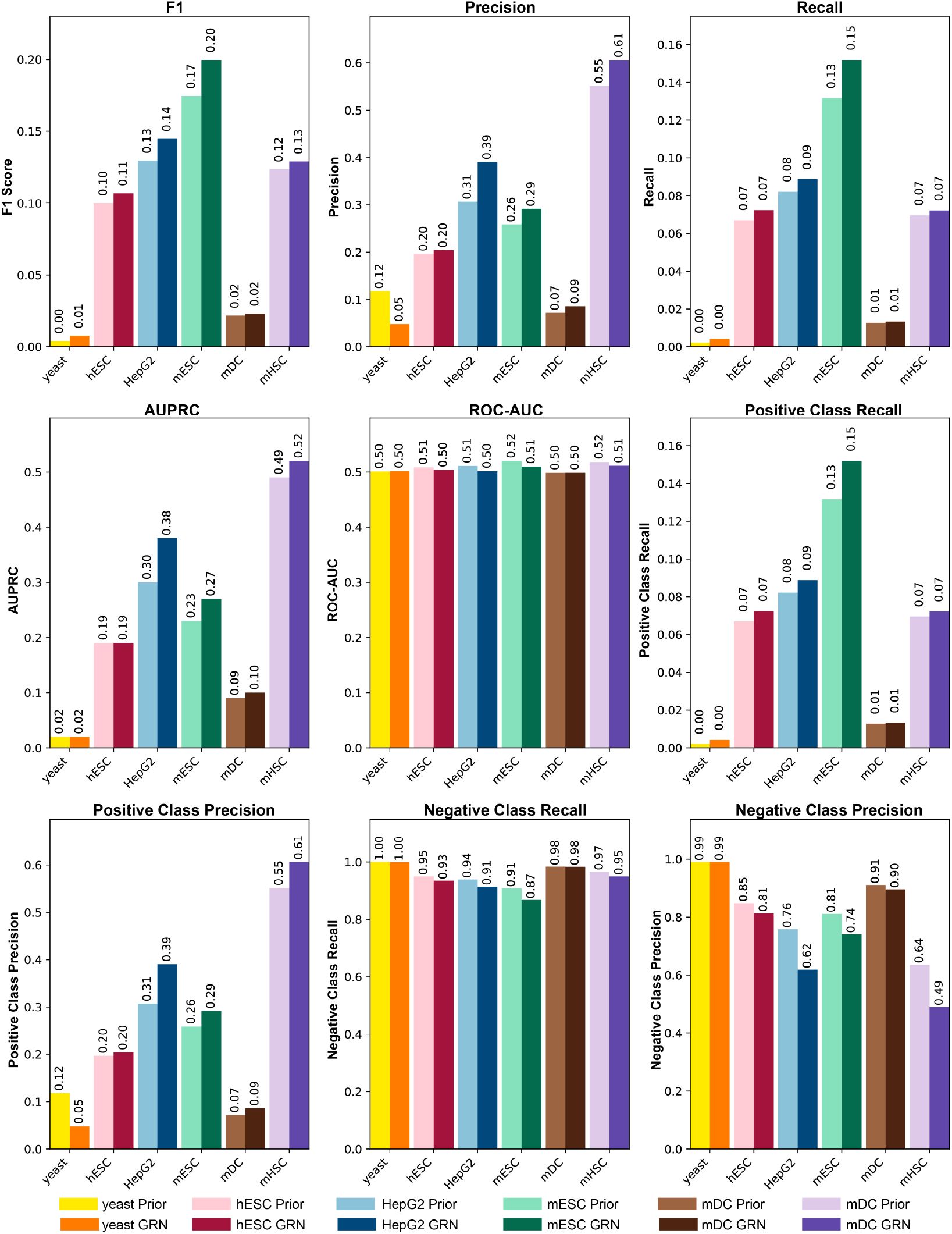
Performance Metrics. Additional performance metrics for GLM-Prior and PMF-GRN across six cell lines.

### F Cross-Method Comparison

In order to disentangle the performance contributions provided by each prior knowledge method (GLM-Prior, Inferelator-Prior, and CellOracle baseGRN) from their corresponding GRN inference method (PMF-GRN, Inferelator 3.0, and CellOracle), we provide a cross-method comparison in which each GRN inference algorithm is applied to each prior. No new datasets are introduced in this section. The results from the cross-method comparison can be found in Table 12.

**Table 12:** Cross-method AUPRC comparison for six cell lines, combining three prior knowledge constructions (CellOracle prior, Inferelator-Prior, GLM-Prior) with three GRN inference methods (PMF-GRN, Inferelator, CellOracle).

| Cell Line | CellOracle Prior |  |  | Inferelator-Prior |  |  | GLM-Prior |  |  |
| --- | --- | --- | --- | --- | --- | --- | --- | --- | --- |
|  | PMF | Inf | CO | PMF | Inf | CO | PMF | Inf | CO |
| Yeast | 0.06 | 0.06 | 0.06 | 0.03 | 0.04 | 0.03 | 0.02 | 0.03 | 0.03 |
| hESC | 0.17 | 0.18 | 0.17 | 0.19 | 0.20 | 0.19 | 0.19 | 0.17 | 0.19 |
| HepG2 | 0.40 | 0.34 | 0.38 | 0.36 | 0.36 | 0.33 | 0.38 | 0.34 | 0.31 |
| mESC | 0.17 | 0.25 | 0.37 | 0.25 | 0.25 | 0.23 | 0.27 | 0.27 | 0.24 |
| mDC | 0.07 | 0.11 | 0.09 | 0.10 | 0.09 | 0.08 | 0.10 | 0.10 | 0.07 |
| mHSC | 0.34 | 0.42 | 0.39 | 0.41 | 0.42 | 0.35 | 0.52 | 0.42 | 0.36 |

